# The connectional diaschisis and normalization of cortical language network dynamics after basal ganglia and thalamus stroke

**DOI:** 10.1101/2024.07.29.605538

**Authors:** Qingwen Chen, Xiaolin Guo, Tao Zhong, Junjie Yang, Xiaowei Gao, Zhe Hu, Junjing Li, Jiaxuan Liu, Yaling Wang, Zhiheng Qu, Wanchun Li, Zhongqi Li, Wanjing Li, Yien Huang, Jiali Chen, Hao Wen, Ye Zhang, Binke Yuan, Han Gao

**Affiliations:** Department of Neurosurgery, The Affiliated Qingyuan Hospital (Qingyuan People’s Hospital), Guangzhou Medical University, Qingyuan, China; Department of Neurosurgery, The First Affiliated Hospital of Guangdong Pharmaceutical University, Guangzhou, China; Key Laboratory of Brain, Cognition and Education Sciences, Ministry of Education, China: Institute for Brain Research and Rehabilitation, South China Normal University, Guangzhou, China; Centre for Cognition and Brain Disorders, Hangzhou Normal University, NO.2318, Yuhangtang Rd, Yuhang District, Hangzhou 311121, China; Philosophy and Social Science Laboratory of Reading and Development in Children and Adolescents (South China Normal University), Ministry of Education, China

**Keywords:** Subcortical stroke, cortico-subcortical interaction, dynamic functional connectivity, dynamic conditional correlation

## Abstract

Stroke affecting the basal ganglia and thalamus can lead to language deficits. In addition to the lesion’s direct impact on language processing, connectional diaschisis involving cortical-subcortical interactions also plays a critical role. This study investigated connectional diaschisis using the “dynamic meta-networking framework of language” in patients with basal ganglia and thalamus stroke, analyzing longitudinal resting-state fMRI data collected at 2 weeks (n = 32), 3 months (n = 19), and one year post-stroke (n = 23). As expected, we observed dynamic cortico-subcortical interactions between cortical language regions and subcortical regions in healthy controls (HC, n = 25). The cortical language network exhibited dynamic domain-segregation patterns in HCs, severely disrupted in the acute phase following stroke. The connectional diaschisis manifested as dual effects characterized by both hypo- and hyper-connectivity, which positively and negatively correlated with language deficits, respectively. State-specific changes in nodal and topological properties were also identified. Throughout language recovery, cortical language network dynamics gradually normalized toward sub-optimal domain-segregation patterns, accompanied by the normalization of nodal and topological properties. These findings underscore the crucial role of cortico-subcortical interactions in language processing.

## Introduction

Stroke predominantly affects subcortical regions and white matter (Corbetta et al., 2015; Kang, Chalela, Ezzeddine, & Warach, 2003; Wessels et al., 2006; B. Yuan et al., 2017), leading to language deficits when occurring in the dominant hemisphere. In addition to the disconnection of language-related white matter tracts, lesions in the basal ganglia and thalamus also contribute to these deficits (Corbetta et al., 2015; Hwang, Shine, Bruss, Tranel, & Boes, 2021; Marcia Radanovic & Almeida, 2021; M. Radanovic & Mansur, 2017). The direct neuropsychological reason is the involvement of basal ganglia and thalamus in language processing. Although the precise roles of basal ganglia and thalamus are still being investigated, sustained activations in these areas have been observed during syntax, semantics, phonology, and prosody processing (David A Copland & Angwin, 2019; D. A. Copland, Brownsett, Iyer, & Angwin, 2021; Hwang, Shine, Cole, & Sorenson, 2022; Ji et al., 2019; Thibault et al., 2021; Turker, Kuhnke, Eickhoff, Caspers, & Hartwigsen, 2023). Lesion-deficit mapping (Corbetta et al., 2015; Hwang et al., 2021; Rangus, Fritsch, Endres, Udke, & Nolte, 2022) and deep brain stimulation (Parsons, Rogers, Braaten, Woods, & Troster, 2006) have also provided causal evidence. Beyond that, several theoretical frameworks have also been laid out, such as the disconnection hypothesis, cortical deafferentation, and diaschisis (Crosson, 2013; Marcia Radanovic & Almeida, 2021). Undoubtedly, these frameworks underscore the importance of cortical-subcortical interactions in language processing. However, studies investigating how basal ganglia and thalamus strokes disrupt these interactions are limited.

Regions in the subcortical regions recruited for language processing are distributed bilaterally and include the bilateral thalamus, putamen, caudate, and amygdala (see the meta-analytic review in Turker et al., 2023). Connections between these language-related subcortical regions and cortical language regions are part of the cortico-thalamo-cortical and cortico-striato-thalamic loops (Bell & Shine, 2016; Greene et al., 2020; Shine, Lewis, Garrett, & Hwang, 2023). Compared to structural connections, more researches have focused on functional interactions. A cortico-subcortical network partition study found that the cortical language-specific areas (including Broca’s area, Wernicke’s area, 55b) had strong functional connectivity with the bilateral amygdala, caudate, and the left putamen (Ji et al., 2019). Furthermore, the functional connectivity patterns in both cortical and subcortical language network showed high left-lateralized asymmetry. Apart from these core cortical language centers, language processing also highly relies on the orbital frontal cortex, middle and inferior temporal gyrus for semantic and syntax processing (Xu, Lin, Han, He, & Bi, 2016) and ventral part of the somatomotor network for speech processing (Lu et al., 2021; Zhao, Liu, Zhang, Lu, & Wu, 2021). These cortical language- and speech-related areas, which belong to multiple resting-state functional networks (e.g., default model network and somatomotor network) had strong functional connectivity with the thalamus, pallidum, putamen, caudate, and amygdala (Ji et al., 2019). Thanks to these densely cortical-subcortical interactions, the thalamic task-evoked responses (including language tasks) have been shown to be highly predictable to cortical task activity (Hwang et al., 2022).

While anatomical and functional descriptions of cortical-subcortical connectivity provide an initial map to understand the lesion effect of basal ganglia and thalamus strokes, empirical evidence is rare. At the whole-brain level, Favaretto et al. (2022) found that cortical regions were temporally and flexibly synchronizing with either limbic regions (hippocampus/amygdala) or subcortical nuclei (thalamus/basal ganglia) within five temporal-reoccurring states in resting state. Subcortical stroke, which damages white matter connections between basal ganglia/thalamus and cortex, disrupted these spatiotemporal patterns of cortical-subcortical dynamics. Critically, they showed that the recovery of spatiotemporal anomalies from 2 weeks to 1 year after stroke was significantly associated with language and memory recovery. However, several limitations of this study remain to be addressed. Firstly, the study by Favaretto et al. (2022) primarily examined the overall cortico-subcortical dynamic interaction at the whole-brain level without specifically delineating the spatiotemporal patterns between subcortical regions and cortical language areas. Previous dynamic studies have shown that the patterns within a network differ from those observed at whole-brain level (Bonkhoff et al., 2020; B. Yuan, Xie, Wang, et al., 2023). Within-network dynamics, which focus on intra-domain dynamics and segregation, may offer greater insights into cognitive specificity and the relevance to behavior or deficits (Bonkhoff et al., 2020; B. Yuan, Xie, Gong, et al., 2023). Secondly, the study did not report the lesion anatomy used for language analysis. Thus, whether the spatiotemporal anomalies resulted from basal ganglia and thalamus stroke or other anatomical structures is unclear. Thirdly, Favaretto and colleagues only constructed a language recovery prediction model based on low-dimensional features. How stroke disrupted the language network dynamics and the normalization of the spatiotemporal anomalies during recovery remains to be elucidated.

To overcome these limitations, in this study, we screened patients with basal ganglia and thalamus stroke from the public dataset used in Favaretto et al. (2022) and investigated how basal ganglia and thalamus stroke disrupted the network dynamics within the cortical language network (connectional diaschisis). Specifically, we adopted our recently proposed “dynamic meta-networking framework of language” to investigate the connectional diaschisis resulting from basal ganglia and thalamus stroke. It is a theoretical framework of language network dynamics in resting state. It includes four temporal-reoccurring states with distinct connectivity patterns, hub distribution, structural underpins, and cognitive relevance (B. Yuan, Xie, Wang, et al., 2023). These four states formed a dynamic “meta-networking” framework of language. Meta-networking refers to a network of networks, which is the key aspect of the “meta-networking” theory of cerebral functions (Herbet & Duffau, 2020). This theory holds that complex cognition and behaviors (e.g., language) arise from the spatiotemporal integration of distributed but relatively specialized sub-networks. The dynamic “meta-networking” framework of language captures the domain-specific nature of language processing, align with the neurobiology of the dual-stream model of speech and language processing (Duffau, Moritz-Gasser, & Mandonnet, 2014; Hickok & Poeppel, 2007), and has been shown to have cognitive and clinical relevance (B. Yuan, Xie, Gong, et al., 2023; B. Yuan, Xie, Wang, et al., 2023). Furthermore, Favaretto et al. (2022) employed the sliding-window method to construct dynamic networks, which have been shown to have several methodological limitations (Lindquist, Xu, Nebel, & Caffo, 2014; Thompson, Richter, Plaven-Sigray, & Fransson, 2018; Xie et al., 2019). Therefore, in this study, we employed the dynamic conditional correlation (DCC) approach (Lindquist et al., 2014) to construct the frame-wise connectivity matrix. DCC has been shown to outperform the standard sliding-window approach in tracking the network dynamics (Choe et al., 2017; Lindquist et al., 2014).

## Materials and Methods

The stroke dataset we used is part of the Washington Stroke Cohort, which has been publicly available at https://cnda.wustl.edu/data/projects/CCIR_00299. The dataset includes longitudinal structural imaging, resting-state fMRI, and neuropsychological testing scores at 2 weeks, 3 months, and one year after stroke. Written informed consent was obtained from all participants in accordance with the Declaration of Helsinki and procedures established by the Washington University in Saint Louis Institutional Review Board. All aspects of this study were approved by the Washington University School of Medicine (WUSM) Internal Review Board.

### Patients

The whole dataset comprised 132 patients with first symptomatic stroke, whose data have been previously published using approaches different from those employed in this study (Corbetta et al., 2015; Favaretto et al., 2022; Ramsey et al., 2016; Ramsey et al., 2017; Salvalaggio, De Filippo De Grazia, Zorzi, Thiebaut de Schotten, & Corbetta, 2020; Siegel et al., 2016; Siegel et al., 2018; Silvestri et al., 2022). The inclusion and exclusion criteria were detailed in Corbetta et al. (2015). We screened patients with basal ganglia and thalamus stroke based on the lesion masks (by neurosurgeons of C.Q and Z.T).

Ultimately, 32 patients at 2 weeks, 19 patients at 3 months, and 23 patients at one year after stroke met the inclusion criteria. Additionally, twenty-five age-, sex- and education-matched healthy controls (HCs) were recruited.

### MRI imaging

MRI imaging data was acquired with a Siemens 3T Tim-Trio scanner at the School of Medicine of Washington University in St. Louis. Structural scans consisted of (1) a sagittal MPRAGE T1-weighted image, TR=1950 msec, TE=2.26 msec, flip angle=9 deg, voxel size=1.0 x 1.0 x 1.0 mm, slice thickness = 1.00 mm; (2) a transverse turbo spin-echo T2-weighted image, TR=2500 msec, TE=435 msec, voxel-size=1.0 x 1.0 x 1.0 mm, slice thickness = 1.00 mm; and (3) a sagittal FLAIR, fluid-attenuated inversion recovery, TR=7500 msec, TE=326 msec, voxel-size=1.5 x 1.5 x 1.5 mm, Slice thickness = 1.50 mm. Resting-state functional scans were acquired with a gradient echo EPI sequence (TR = 2000 ms, TE = 27 ms, 32 contiguous 4 mm slices, 4 × 4 mm in-plane resolution) during which participants were instructed to fixate on a small white cross-centered on a screen with a black background in a low luminance environment. Six to eight resting state (RS) fMRI runs, each including 128 volumes (30 min total), were acquired.

### Data preprocessing

The resting-state fMRI data were preprocessed in the following steps: 1) delete the first 10 volumes; 2) slice timing; 3) head motion correction (< 3 mm or 3 degrees); 4) coregister; 5) T1 segmentation and normalization into the MNI space using DARTEL; 6) functional normalization using the deformation field of T1 images; 7) smoothing using a Gaussian kernel (full-width-at-half-maximum = 6 mm); 8) Linear detrend; 9) Nuisance signals regression (24 motion parameters, including the x, y, z translations, and rotations (6 parameters), plus their temporal derivatives (6 parameters) and the quadratic terms of 12 parameters, and white matter/cerebrospinal fluid/global mean time courses); 10) Temporal band-pass filtering (0.01–0.1 Hz).

### The cortical language network definition

To investigate the connetional diaschisis, we defined a putative cortical language network according to the Human Brainnetome Atlas (Fan et al., 2016). We extracted all parcels that exhibited significant activation in language-related behavioral domains (i.e., language or speech) or paradigm classes (e.g., semantic, word generation, reading, or comprehension). Thirty-three cortical parcels in the left hemisphere and 15 cortical parcels in the right hemisphere were selected. Their extents are highly similar to those identified by other researchers (Lipkin et al., 2022; Vigneau et al., 2006). Considering bilateral language processing (Hodgson, Lambon Ralph, & Jackson, 2021; Siegel et al., 2016; Turker et al., 2023; Vigneau et al., 2011) and the recruitment of right hemisphere language regions for recovery, 19 parcels of homologs in the right hemisphere and 1 parcel of homologs in the left hemisphere were also selected. Altogether, a symmetric language network (BNL68, Supplementary Figure 1) including 68 cortical ROIs (34 ROIs in each hemisphere) was defined (Binke Yuan et al., 2022), including the superior, middle, and inferior frontal gyrus (SFG, MFG, and IFG, respectively), the ventral parts of the precentral gyrus (PrG) and postcentral gyrus (PoG), the STG, MTG and inferior temporal gyrus (ITG), the fusiform gyrus (FuG), the Para hippocampus gyrus (PhG) and the posterior superior temporal sulcus (pSTS). The coordinates (in Montreal Neurologic Institute space) and meta results of each parcel are summarized in Supplementary Table 1.

### A cortico-subcortical atlas for dynamic cortico-subcortical analysis

To construct the cortical-subcortical network, we selected subcortical parcels (n = 36), including 4 subregions for the amygdala, 4 subregions for the hippocampus, 12 subregions for basal ganglia, and 16 subregions for thalamus, from the Human Brainnetome Atlas were selected. We included all subcortical regions due to their widespread involvement in language processing. In total, 104 cortical and subcortical regions were selected for cortical-subcortical language network analysis.

It should be noted that due to damage in the subcortical regions of the patient group, the dynamic analysis between the cortical language network and subcortical regions was conducted only in HCs. Given the limited sample size of HCs (only 25 subjects), we utilized a publicly available dataset comprising 192 healthy college students to validate the robustness of our findings. This dataset has been demonstrated with high imaging quality and is the same dataset used for proposing the dynamic meta-networking framework of language model (B. Yuan, Xie, Wang, et al., 2023). The 192 subjects (118 females; aged 18–26 years old, mean ± STD = 21.17 ± 1.83) were a part of the 1000 Functional Connectomes Project (Beijing data, available at http://fcon_1000.projects.nitrc.org). For each subject, R-fMRI data and T1-weighted images were acquired using a SIEMENS TRIO 3-Tesla scanner. The R-fMRI data were collected with the following parameters: repetition time = 2s, 33 axial slices, and 225 frames. The preprocessing procedures were the same as those of HCs.

### Frame-wise time-varying network construction using DCC

To identify the temporal reoccurring states of two types of networks, the dynamic conditional correlation (DCC) approach was adopted to construct the framewise time-varying language network (https://github.com/caWApab/Lindquist_Dynamic_Correlation/tree/master/DCC_tool box).

DCC is a variant of the multivariate GARCH (generalized autoregressive conditional heteroscedastic) model (Lebo & Box-Steffensmeier, 2008) (<u>Engle, 1982</u>; <u>Lebo & BoxlJSteffensmeier, 2008</u>), which has been shown to be particularly effective for estimating both time-varying variances and correlations. GARCH models express the conditional variance of a single time series at time *t* as a linear combination of the past values of the conditional variance and of the squared process itself. All the parameters of DCC are estimated through quasi-maximum likelihood methods and require no ad hoc parameter settings.

The DCC algorithm consists of two steps. To illustrate, let us assume that there is a pair of time series from two ROIs, x_t_ and y_t_. In the first step, standardized residuals of each time series are estimated using a univariate GARCH (1,1) process. In the second step, an exponentially weighted moving average (EWMA) window is applied to the standardized residuals to compute a non-normalized version of the time-varying correlation matrix between x_t_ and y_t_. The mathematical expressions of the GARCH (1,1) model, DCC model, and EWMA, and the estimations of the model parameters were provided by <u>Lindquist et al.</u> (<u>2014</u>).

K-means clustering was adopted to decompose the dFC matrices into several reoccurring connectivity states. The optimal number of clusters *k* was estimated based on elbow criterion, i.e., the ratio between the within-cluster distance to between-cluster distances (<u>Allen et al., 2014</u>). L1 distance function (‘Manhattan distance’) was implemented to assess the point-to-centroid distance. Each time window was finally assigned to one of these connectivity states.

### Topological properties of dFC states

A state-specific subject connectivity matrix was estimated by calculating the median of all connectivity matrices that were assigned to the same state label (Damaraju et al., 2014) (<u>Damaraju et al., 2014</u>). Before calculation, a correlation threshold (*r* > 0.2) was used to eliminate weak connections that might have arisen from noise. Due to the nature of negative connections being debatable (Murphy & Fox, 2017), only positive connections were considered in this work. In the current study, we used a weighted network rather than a binarized network to conserve all connectivity information.

*<u>Global topological metrics</u>*: for each state-specific matrix, three network topological properties were calculated: 1) Total connectivity strength. The FC strength of a network was calculated by summing functional connectivity strengths of all suprathreshold connections into one value (Nelson, Bassett, Camchong, Bullmore, & Lim, 2017); 2) Global network efficiency (gE). The gE reflects the capability for parallel information transfer and functional integration. It is defined as the average of the inverses for all weighted shortest path lengths (the minimal number of edges that one node must traverse to reach another) in the thresholded matrix (Rubinov & Sporns, 2010); 3) Local network efficiency (lE). The lE reflects the relative functional segregation and is defined as the average of the global efficiency of each node’s neighborhood sub-graph.

*<u>Nodal topological metrics</u>*: for each node, nodal strength, gE, and lE were calculated. We also calculated betweenness centrality (BC), an index indicating whether a particular node lies on the shortest paths between all pairs of nodes in the network.

As a complementary analysis, we also calculated the static functional connectivity (sFC) matrix using the Pearson correlation coefficient. We then calculated the three topological properties of sFC and investigated the alterations in the two patient groups.

### Functional relevance of state-dependent hub distributions

To assess the functional relevance of hubs in each state, we performed term-based meta-analyses on the platform Neurosynth (https://www.neurosynth.org/). Neurosynth performs an automated selection of studies based on the predefined term. It divides the entire database of coordinates into two sets: those that occur in articles containing a particular term and those that do not. Then it performs meta-analysis using forward inference maps, which reflect the likelihood that a voxel will activate if a study uses a particular term. To verify the functional specificity, five speech perception, inner speech, phonological processing, speech production, and semantic processing, were analyzed. The functional relevance was assessed by calculating the dice coefficients between binary images of hub nodes and meta results (dice = 2* (hub * meta)/(hub + meta)). Considering the left-lateralized activations of meta results, the dice coefficients were calculated in the left hemisphere.

### Statistical analysis

Age and education data were analyzed by two-sample t-tests. Gender, lesion type, race, smoke, high blood pressure, and diabetes were analyzed by Pearson’s chi-square test. The language tests and PCA scores between HCs and patients were analyzed by a linear mixed effect model.

The changes in functional connectivity and nodal and global topological properties were analyzed by two-sample t-tests. The results of edge strength were corrected using network-based statistics (NBS) (Zalesky, Fornito, & Bullmore, 2010) with an edge *p*-value of 0.05 and a component *p*-value of 0.05. The results of nodal and global topological properties were corrected using False Discovery Rate (FDR) with a corrected *p-value* of 0.05.

The associations between edge, nodal, and global topological properties and the patients’ language scores were assessed by calculating the partial Pearson correlation coefficient after regressing out age, gender, lesion type (including ischemic and hemorrhagic), race, and education. The results between edge strength and language scores were corrected using NBS with a component *p*-value of 0.05.

## Results

### Lesion anatomy, language deficits, and recovery

Patients with circumscribed lesions in the thalamus, pallidum, caudate, amygdala, and putamen, with the highest overlap in the bilateral limb of the internal capsule (Figure 1). Whiter matter disconnections were also observed in the corticobulbar tract (ID = 7), the occipital part of the corticopontine tract (ID = 30), the corticospinal tract (ID = 31), the dentatorubrothalamic tract (ID = 50), the frontal aslant tract (ID = 53), the medial lemniscus (ID = 64), and the superior part of the thalamic radiation (ID = 85).

**Figure 1.**
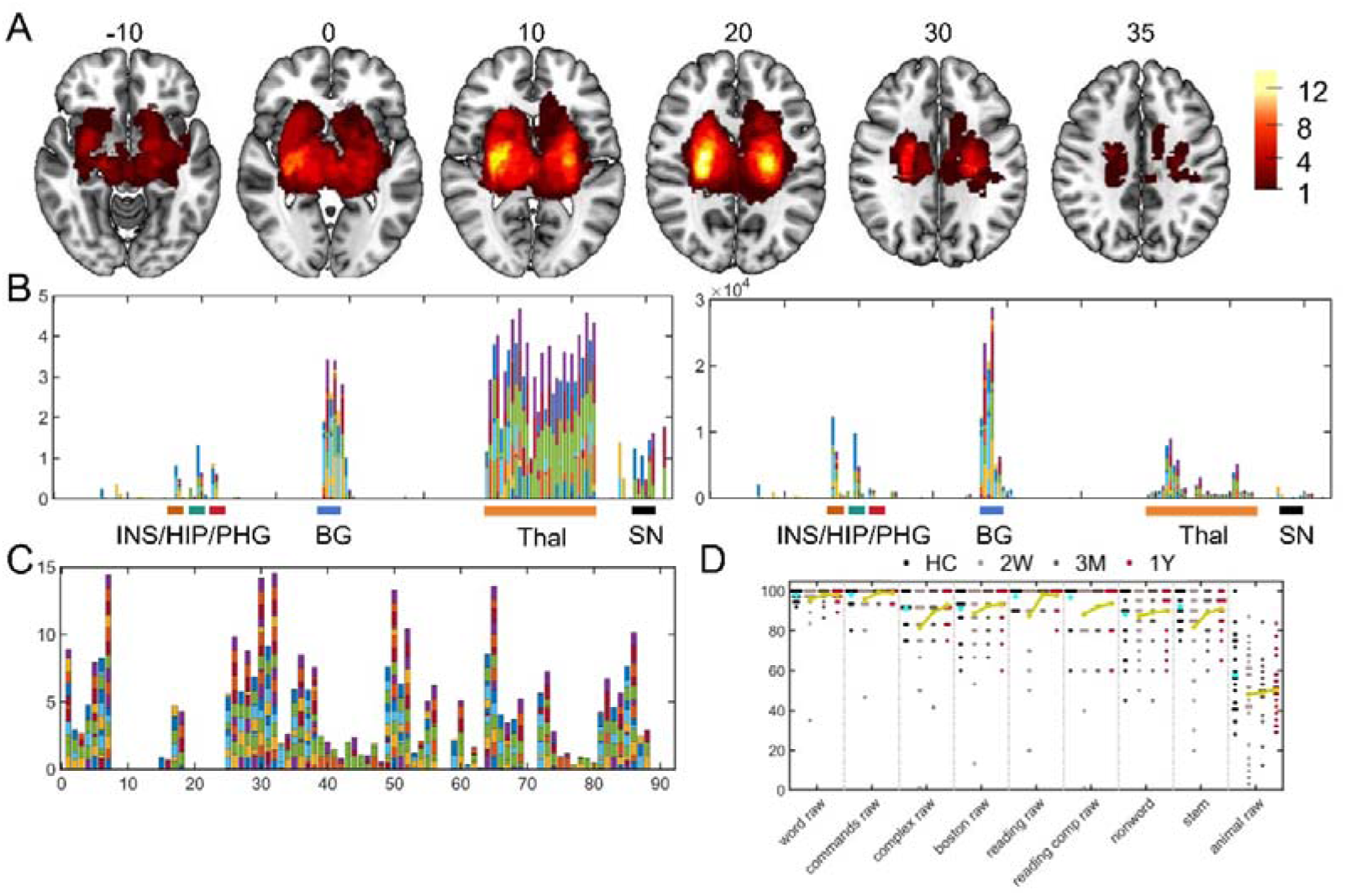
Lesion anatomy, white matter disconnection, and language deficits and recovery. A: the lesion overlap map in 2 weeks after stroke. The color bar represents the number of patients with a lesion on a specific voxel. B: the lesion percentages (left) and total lesion voxel numbers (right) for each subcortical region. The anatomy of the subcortical region was from the Brainnetome atlas (Fan et al., 2016). C: the amount of white matter disconnections for each patient. A group-averaged tractography atlas with 91 white matter tracks, i.e., the HCP-1065 atlas (Yeh, 2022), was adopted to quantify the white matter disconnections. D: language tests for HCs and patients.

**Figure 2.**
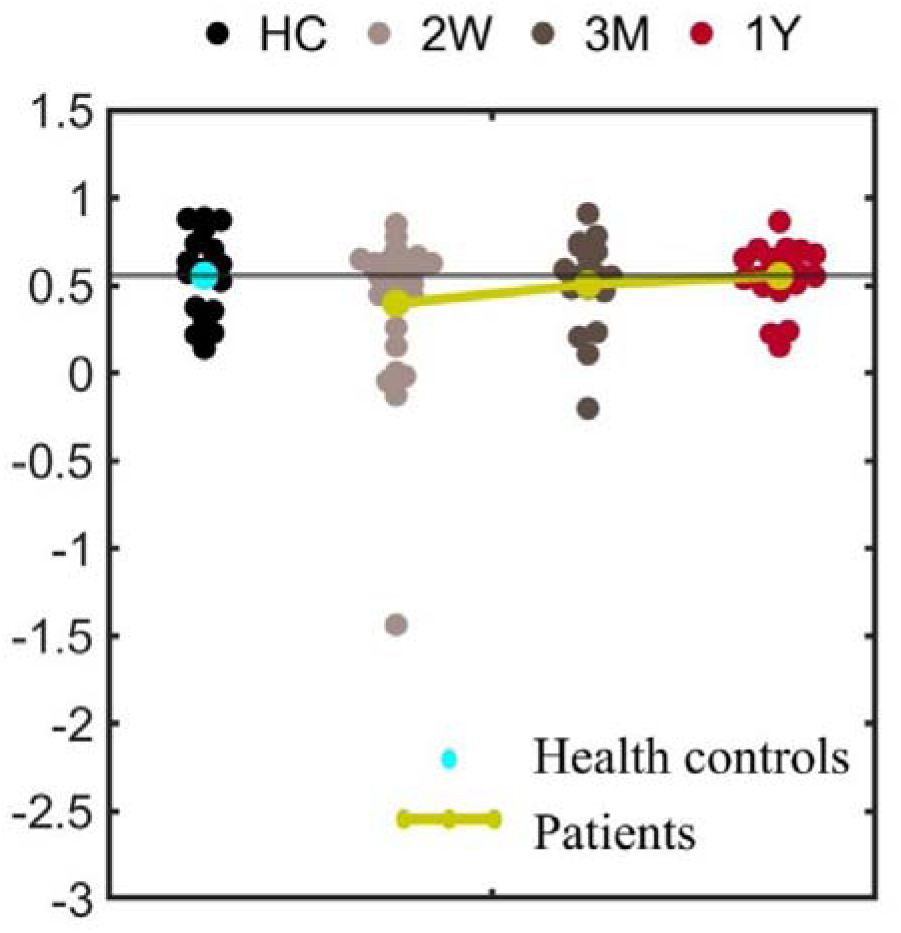
The PCA-based language deficits in acute phase and recovery in 3 months and 1 year after stroke. Linear mixed effect model analyses showed a significant group effect (*P* = 0.006) and post-stroke time effect (*P* = 0.015).

Table 1 summarizes the demographic and clinical information. There was no significant between-group difference for age, sex composition, race, and education. Patients showed significant higher high blood pressure compared with HCs. Language deficits were observed two weeks after stroke, which resulted from severe deficits in subtests of complex ideational material, oral reading of sentences, stem completion and comprehension of oral reading of sentences (Table 2). These language deficits recovered the most within three months after stroke.

**Table 1.**
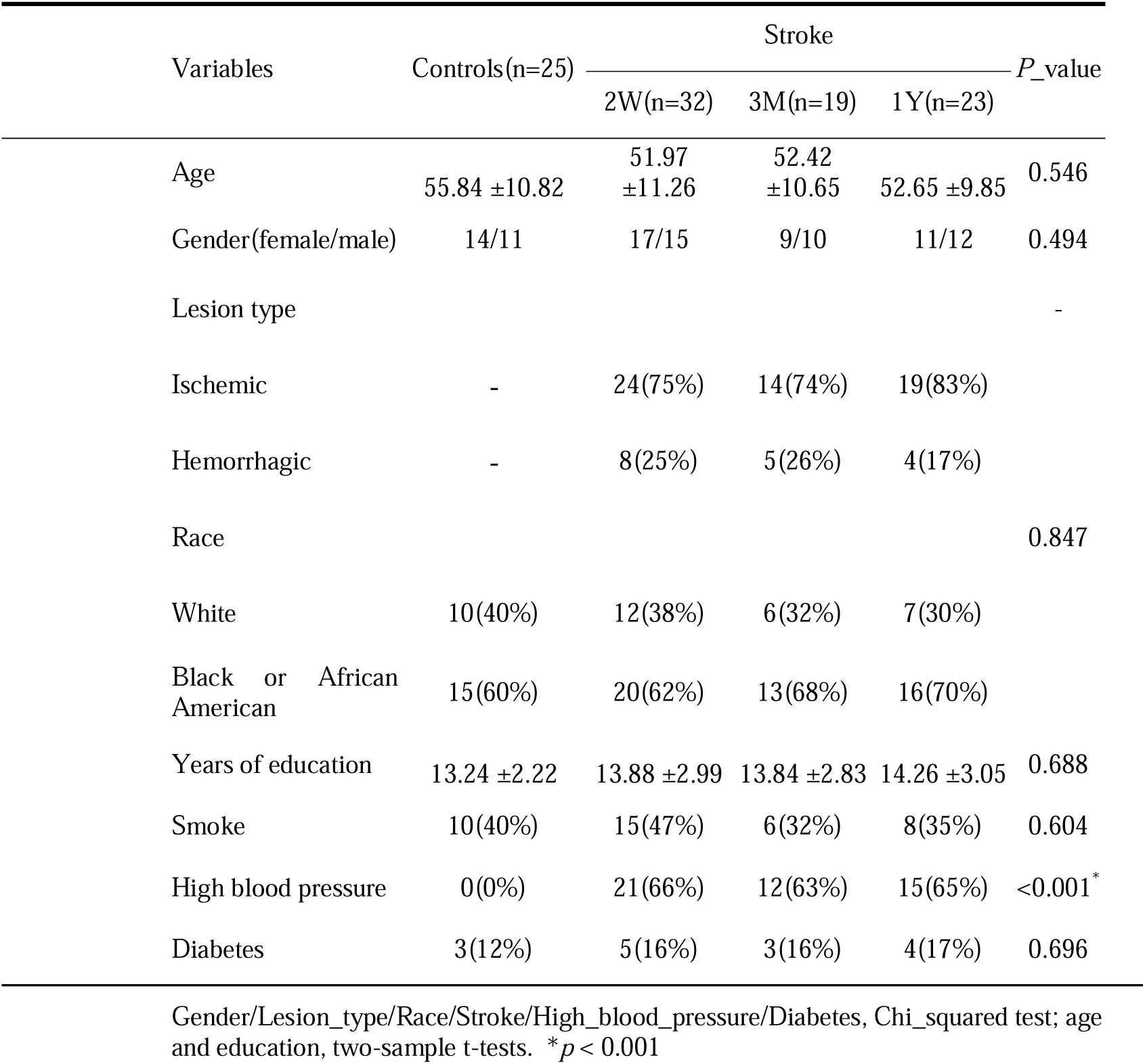
Demographic and clinical information of stroke patients and healthy controls.

| Variables | Controls(n=25) | Stroke |  |  | P_value |
| --- | --- | --- | --- | --- | --- |
|  |  | 2W(n=32) | 3M(n=19) | 1Y(n=23) |  |
| Age | 55.84 ±10.82 | 51.97 ±11.26 | 52.42 ±10.65 | 52.65 ±9.85 | 0.546 |
| Gender(female/male) | 14/11 | 17/15 | 9/10 | 11/12 | 0.494 |
| Lesion type |  |  |  |  | - |
| Ischemic | - | 24(75%) | 14(74%) | 19(83%) |  |
| Hemorrhagic | - | 8(25%) | 5(26%) | 4(17%) |  |
| Race |  |  |  |  | 0.847 |
| White | 10(40%) | 12(38%) | 6(32%) | 7(30%) |  |
| Black or African American | 15(60%) | 20(62%) | 13(68%) | 16(70%) |  |
| Years of education | 13.24 ±2.22 | 13.88 ±2.99 | 13.84 ±2.83 | 14.26 ±3.05 | 0.688 |
| Smoke | 10(40%) | 15(47%) | 6(32%) | 8(35%) | 0.604 |
| High blood pressure | 0(0%) | 21(66%) | 12(63%) | 15(65%) | <0.001* |
| Diabetes | 3(12%) | 5(16%) | 3(16%) | 4(17%) | 0.696 |

Gender/Lesion\_type/Race/Stroke/High\_blood\_pressure/Diabetes, Chi\_squared test; age and education, two-sample t-tests. \* $p < 0.001$

**Table 2.**
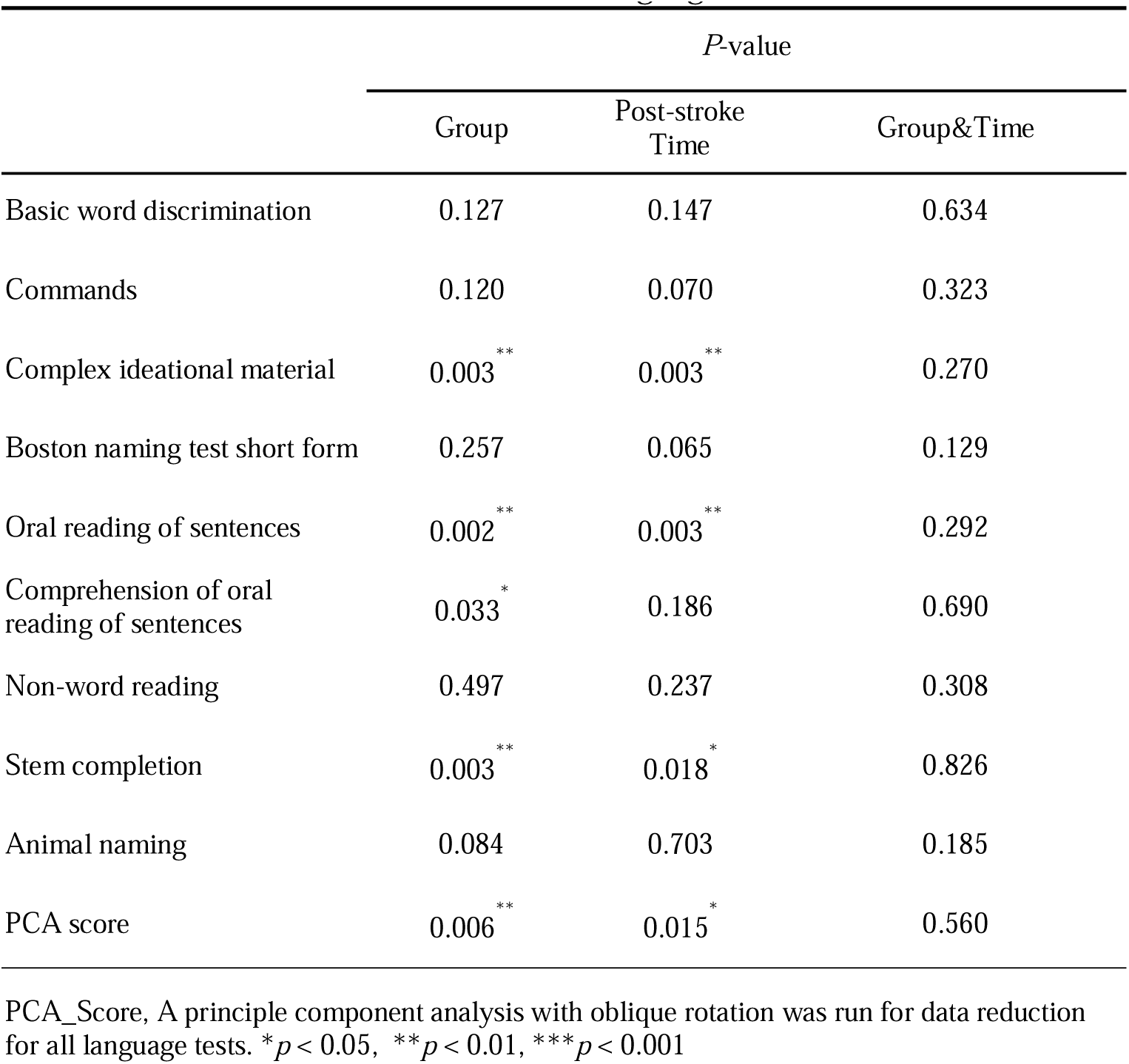
linear mixed effect results of language tests.

|  | *P*-value |  |  |
| --- | --- | --- | --- |
|  | Group | Post-stroke Time | Group&Time |
| Basic word discrimination | 0.127 | 0.147 | 0.634 |
| Commands | 0.120 | 0.070 | 0.323 |
| Complex ideational material | 0.003<sup>**</sup> | 0.003<sup>**</sup> | 0.270 |
| Boston naming test short form | 0.257 | 0.065 | 0.129 |
| Oral reading of sentences | 0.002<sup>**</sup> | 0.003<sup>**</sup> | 0.292 |
| Comprehension of oral reading of sentences | 0.033<sup>*</sup> | 0.186 | 0.690 |
| Non-word reading | 0.497 | 0.237 | 0.308 |
| Stem completion | 0.003<sup>**</sup> | 0.018<sup>*</sup> | 0.826 |
| Animal naming | 0.084 | 0.703 | 0.185 |
| PCA score | 0.006<sup>**</sup> | 0.015<sup>*</sup> | 0.560 |

PCA\_Score, A principle component analysis with oblique rotation was run for data reduction for all language tests. <sup>\*</sup> $p < 0.05$ , <sup>\*\*</sup> $p < 0.01$ , <sup>\*\*\*</sup> $p < 0.001$

### The cortical language network dynamics and cortico-subcortical dynamics

#### The domain-segregation cortical language network dynamics

In HCs, four temporal reoccurring states with distinct connectivity patterns and hub distribution were identified (Figure 3), which aligned with the “dynamic meta-networking framework of language.” State 1 was characterized by moderate to high positive connectivity between nodes in the bilateral PrG, PoG, and STG, but weak or moderate negative connectivity between the prefrontal nodes and temporal nodes. We observed strong positive connectivity among the prefrontal nodes in state 2. State 3 distinguished itself from states 1 and 2 due to strong connectivity among temporal nodes. State 4 showed an overall weak connectivity pattern.

**Figure 3.**
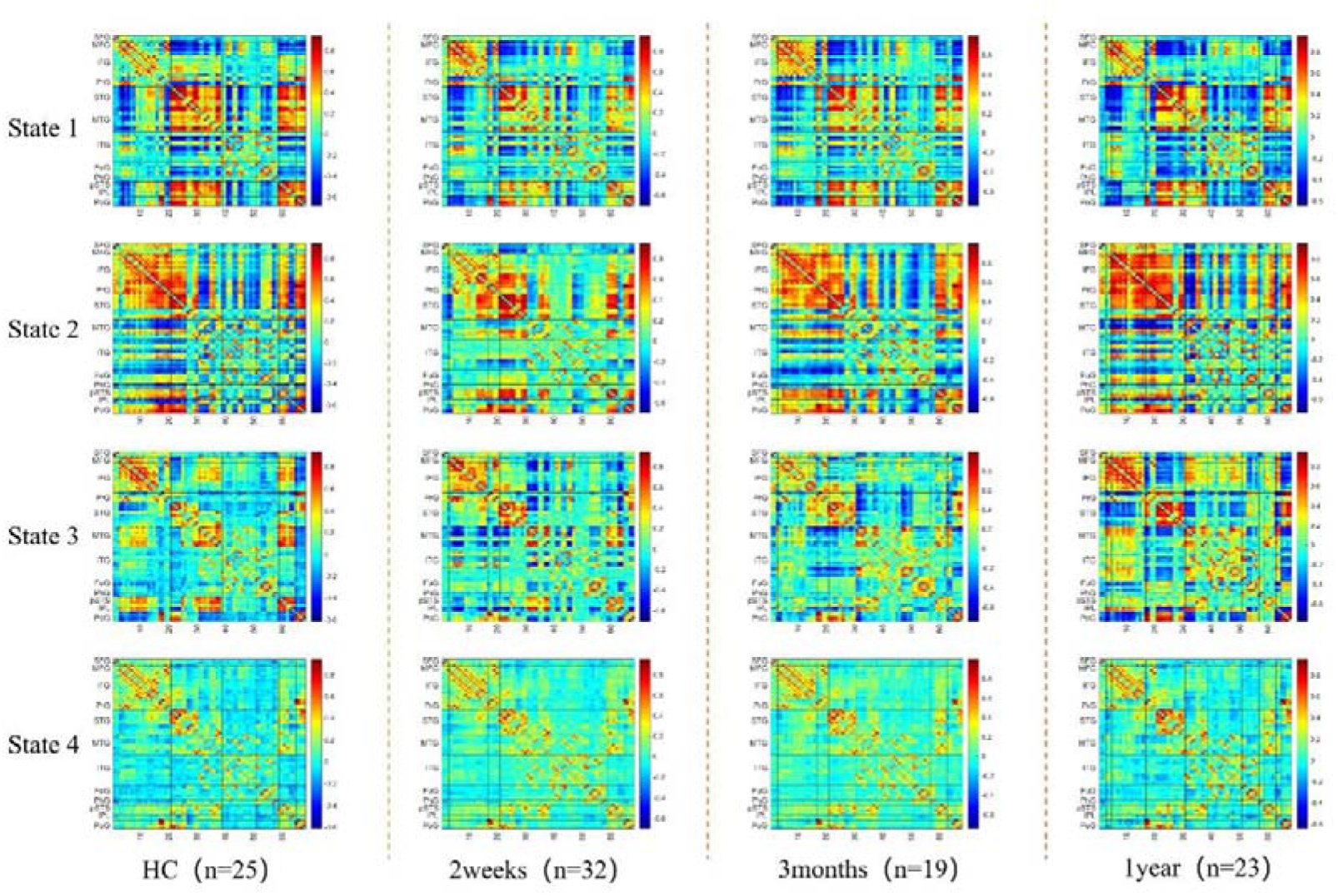
The four temporal reoccurring states in HCs and patients. Warm color denotes positive connections, while code color denotes negative connections.

**Figure 4.**
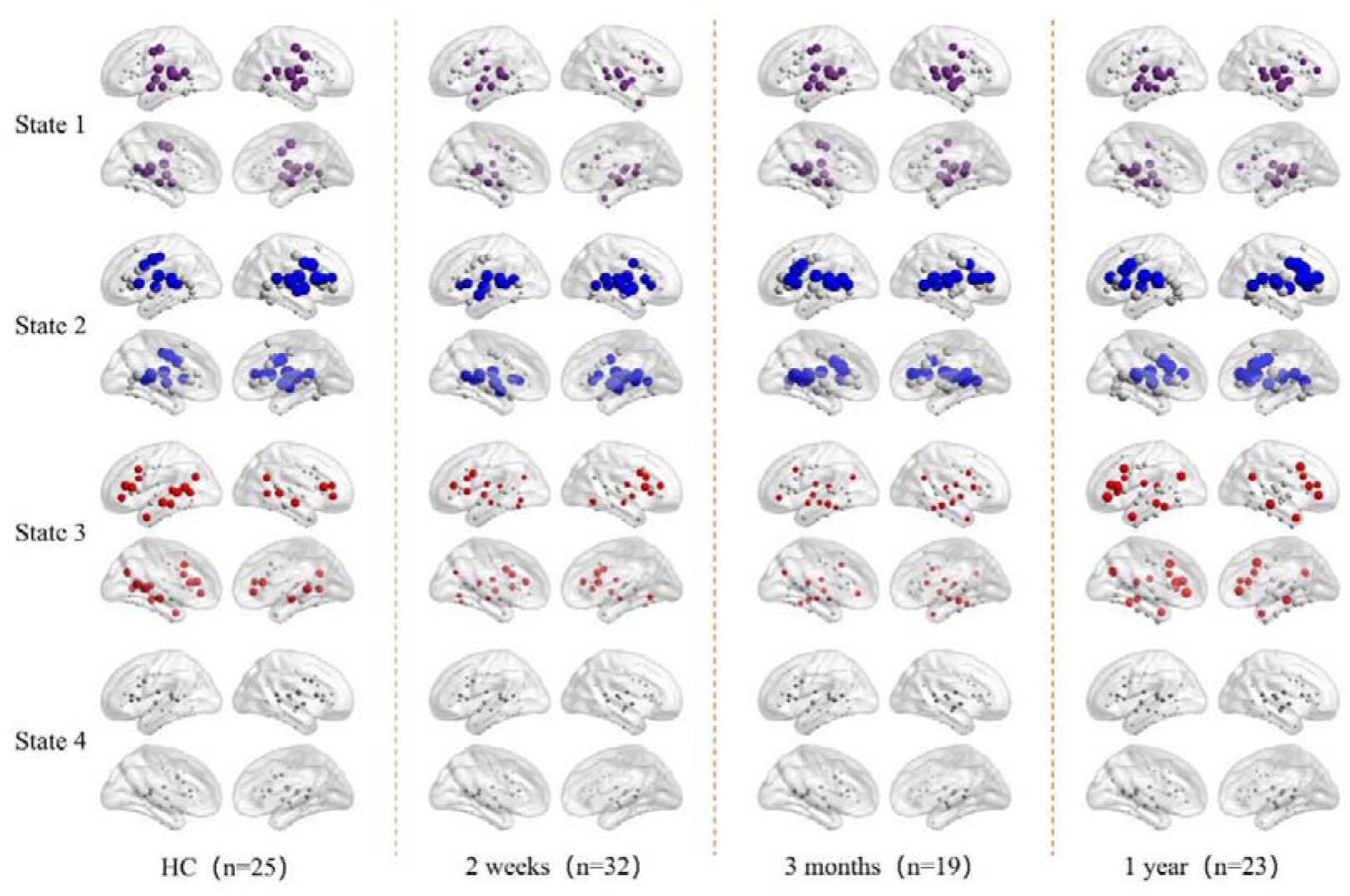
The state-dependent hub distributions. In the first three states, the first 20 nodes with the highest nodal strength were colored and defined as hubs.

The nodal strength distributions of all states were best fitted by the exponentially truncated power-law form (Supplementary Figure 2), which suggested the existence of a small set of highly connected hub nodes. The first 20 nodes with the highest nodal strengths were defined in each group as hubs. In state 1, hubs were mainly distributed in STG. In state 2, hubs were primarily distributed in the prefrontal and posterior temporal cortex. In state 3, hubs were primarily distributed in the temporal cortex, posterior parietal cortex, and IFG. We did not consider the hubs in state 4 because of the weak connectivity strength.

By calculating the spatial similarity between term-based activation and hub distribution, we found that hubs in the four states highly overlapped with activation of speech perception, inner speech, phonological processing, and speech production and partly overlapped with semantic processing (Supplementary Figure 3). Considering the critical role of STG in speech perception and phonological processing, we speculated that hubs in state 1 were mainly implicated in speech perception and phonological processing. Considering the critical role of IFG and the ventral part of PrG in speech production, we speculated that hubs in state 2 were mainly implicated in speech production. Considering the critical role of IPL, ATL, and orbitofrontal cortex in semantic processing, we speculated that hubs in state 3 were mainly implicated in semantic processing. State 4 may be a transition state among these cognition-specific states.

### The cortico-subcortical dynamic interactions

To elucidate the cortico-subcortical dynamic, we calculated the framewise functional connectivity between the 68 cortical language regions and 36 subcortical regions. The cortico-subcortical dynamic also demonstrated four temporal reoccurring states in HCs (Supplementary Figure 4). In the first three states, BG and thalamus consistently exhibit tight connections, but their patterns of connectivity with the cortical language network vary. In State 1, BG and thalamus are positively connected with frontal language areas (IFG and SFG) but negatively connected with regions such as STG, MTG, PrG, and PoG. They show weaker negative connections with regions like ITG, hippocampus, and amygdala. In State 2, BG and thalamus strengthen their connections with the hippocampus and amygdala but weaken connections with other cortical language areas. In State 3, BG and thalamus reduce their connections with IFG but enhance connections with language areas in the sensorimotor network (PrG and PoG). State 4 differs significantly from the first three states, exhibiting overall weaker connections. Connections within the same sub-region of a brain area show moderate positive connections, whereas connections between different brain areas exhibit weak connections. Interestingly, analyzing only the connectivity patterns within the cortical language regions reveals domain-segregation patterns, indicating that while there are functional interactions between cortical language regions and subcortical areas, interactions within the cortical language network itself are relatively independent.

We observed similar cortico-subcortical dynamics in young healthy subjects (see Supplementary Figure 5). The four states showed high spatial similarity across the two groups of healthy subjects (Supplementary Figure 6).

### The disruptions of cortical language network dynamics and recovery over time

Patients’ cortical language network dynamics also exhibited four temporal reoccurring states (Figure 3). Two weeks after stroke, although the cortical language network dynamics were severely disrupted (Figure 5 and Supplementary Figure 7), nodal strength distributions in the four states were best described by the exponentially truncated power-law form (Supplementary Figure 2). We still defined the first 20 nodes with the highest nodal strength as hubs. Distinctive hub distributions were observed in the first three states compared to HCs. In state 1, hubs were observed in pSTG but with weaker nodal strength. In state 2, connections in IFG were weak, and hubs were only observed in pSTG. In state 3, hubs with weaker nodal strength were observed in IFG, pSTG and AG. During language recovery, the functional connectivity recovered, and the domain-segregation hub distributions reemerged, especially for state 2 and 3.

**Figure 5.**
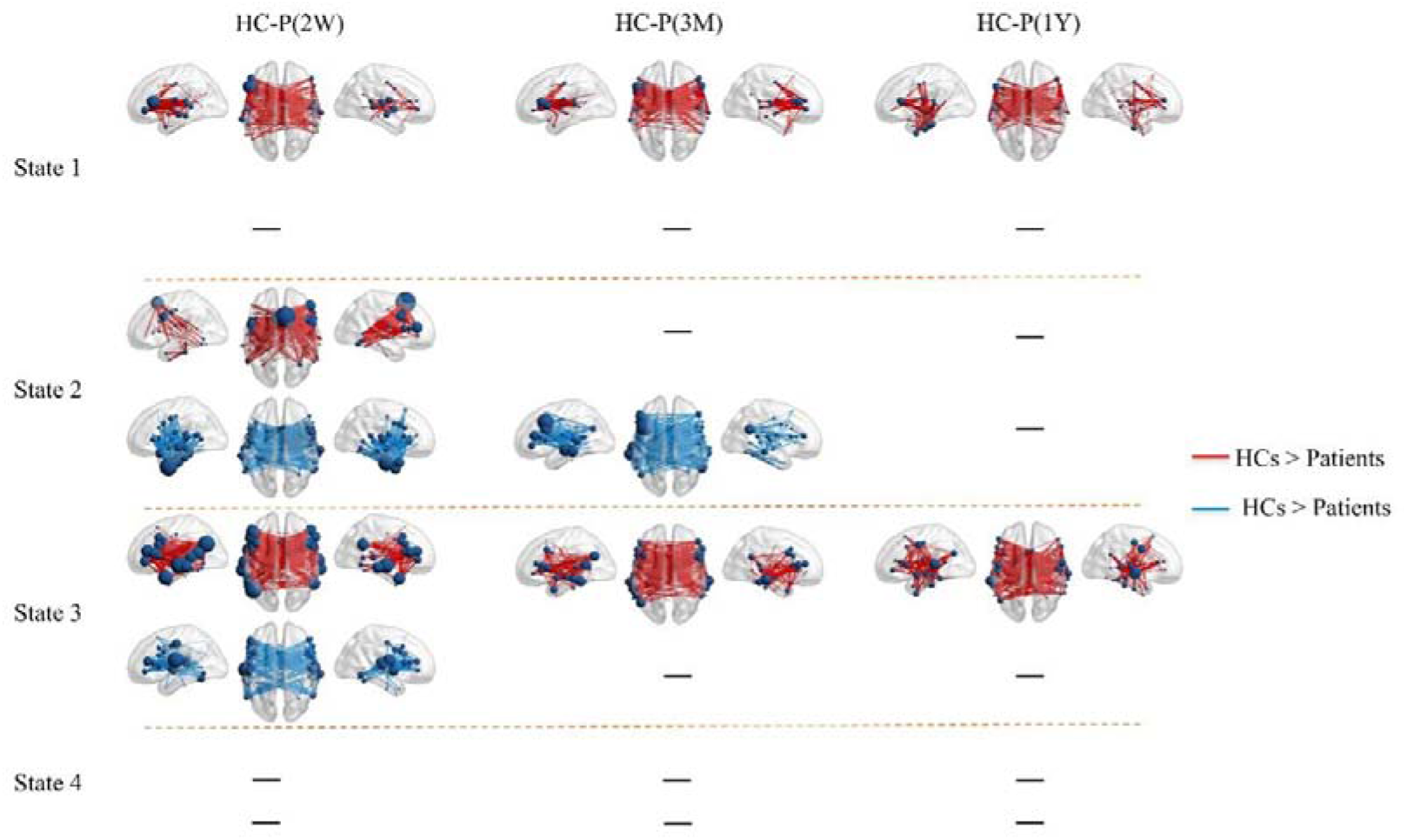
The state-dependent network disruptions in 2 weeks after stroke and normalization during language recovery. Edge *p*<.05, component *p*<.05 with NBS correction.

Statistical analyses showed that the functional connectivity of the first three states was severely disrupted at 2 weeks after stroke. Both hypo- and hyper-connectivity were observed (Figure 5). Interhemispheric connections dominate these hypo- and hyper-connectivities. Partial correlation analysis revealed significant positive and negative correlations between hypo- and hyper-connectivity and language performance, respectively, indicating that connectional diaschisis manifests dual effects (Figure 6).

**Figure 6.**
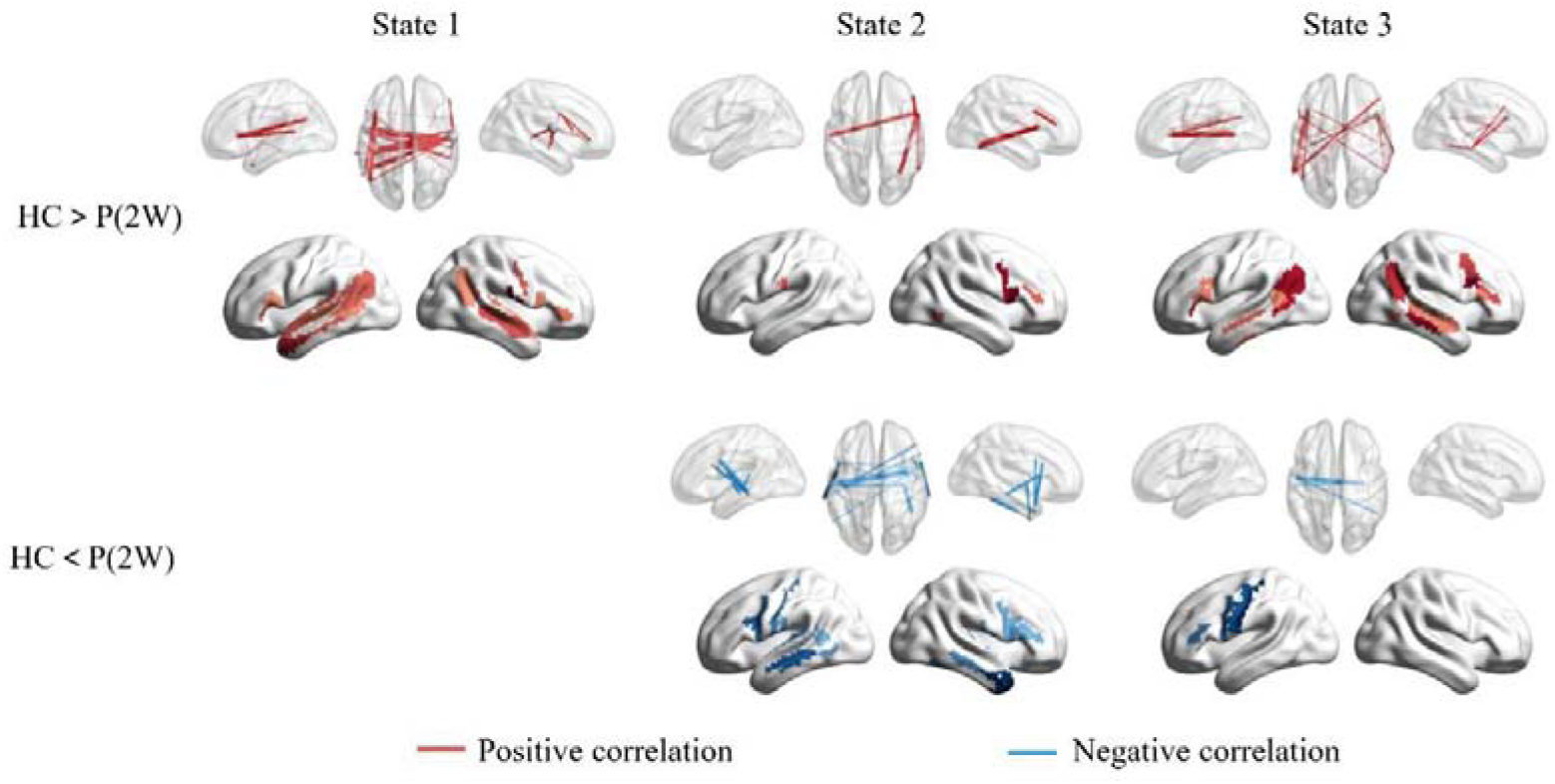
The partial correlation between network disruptions and PCA-based language scores in 2 weeks after stroke. For illustrative purposes, we also sum the significantly correlated edges for each node. Edge *p*<.05, component *p*<.05 with NBS correction.

In state 1, there is a significant reduction in the nodal BC of the left IFG. In state 2, there were significant reductions of nodal DC, gE and lE in the left SFG. Partial correlation analyses revealed positive correlations between nodal gE and PCA-based language score (*R* = 0.34, uncorrected *p* = 0.04) and between nodal lE and PCA-based language score (*R* = 0.55, uncorrected *p* = 0.0004).

In state 3, there is a significant increase in nodal strength and gE of bilateral STG and ITG (Figure 7). The global network efficiency of state 3 also shows a significant increase (uncorrected *p* < 0.01), although it does not withstand FDR correction (Figure 8). No significant correlation between global gE and PCA-based language score were observed.

**Figure 7.**
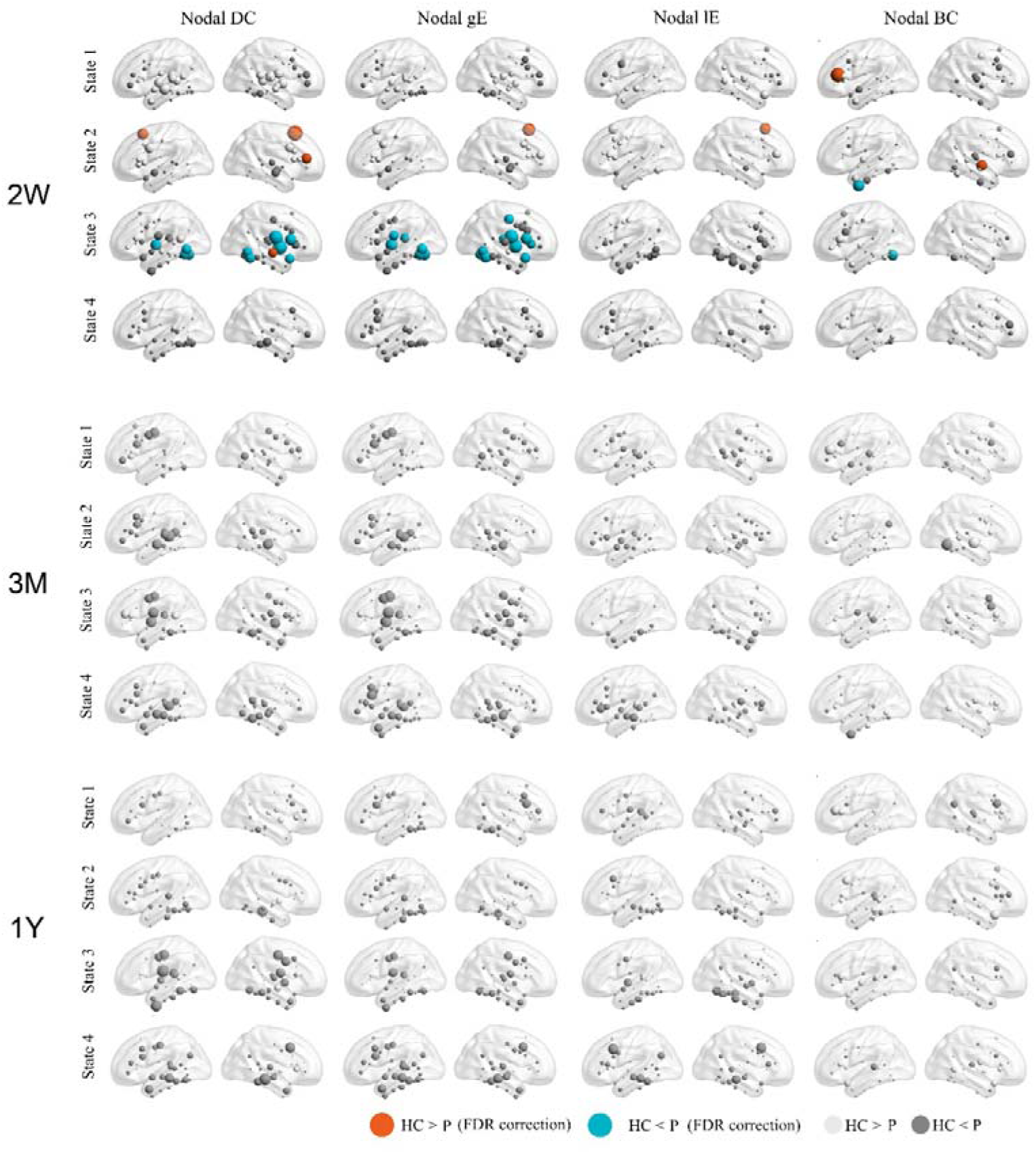
Changes of nodal properties in acute phase and normalization during language recovery. Four nodal properties, the nodal strength, nodal global efficiency (gE), nodal local efficiency (lE), and nodal betweenness centrality (BC) were calculated for each subject median of state dFCs. Between-group statistical comparisons were performed by using -sample t-tests (FDR correction, *p* < 0.05).

**Figure 8.**
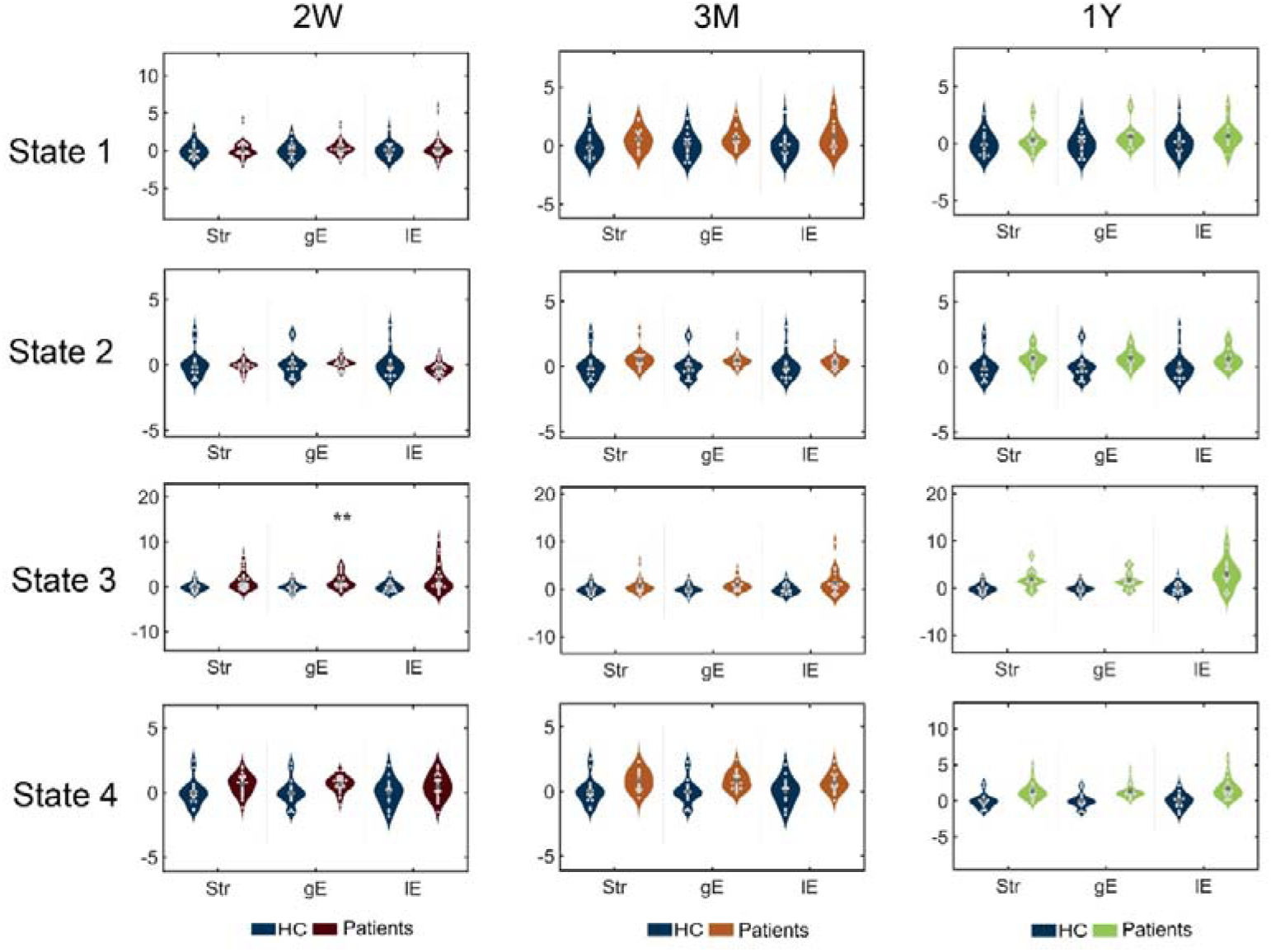
Changes of global properties in 2 weeks after stroke and normalization during language recovery. Increases of network gE were observed in state 3 at 2 weeks after stroke (*p* < 0.01), but did not reach statistical significance (FDR correction with a corrected *p* value 0.05).

In 3 and 12 months after stroke, network disruptions were still observed in states 1 and 3, but the nodal and global topological properties normalized, and no significant difference was observed.

### The properties of sFC analysis

No significant difference in sFC was observed between HCs and patients. Similar functional connectivity patterns were observed for all subjects (Supplementary Figure 8).

## Discussion

Language processing relies on cortico-subcortical interactions, and strokes affecting basal ganglia and thalamus disrupt these interactions, leading to language deficits, a phenomenon termed connectional diaschisis. This study investigated how basal ganglia and thalamus stroke affected the fine-grained cortical language network dynamics. We first constructed a dynamic meta-networking framework of the cortical language network in healthy controls and then investigated how basal ganglia and thalamus stroke disrupted the domain-segregation cortical language network dynamics in the acute phase and recovered within the first year. In the acute phase, cortical language network dynamics were disrupted with state-specific hypo- and hyper-connectivity patterns. The hypo- and hyper-connectivity were positively and negatively correlated with language deficits, respectively, which suggested dual effects. During language recovery, a sub-optimal domain-segregation cortical language network dynamics re-emerged. In healthy subjects, we also observed reliable cortico-subcortical dynamics, which may be network bases of the remote lesion effects. These results refined the connectional diaschisis of basal ganglia and thalamus stroke to the cortical language network and highlighted the importance of cortico-subcortical interaction in language processing.

### The domain-segregation cortical language network dynamics in resting states

The “dynamic meta-networking framework of cortical language network” was first proposed by Yuan et al. (2023). It includes four states with distinctive hub distributions. The meta-analytic results suggest that these hubs are domain-specific. The framework can be regarded as a dynamic and connectional account of speech processing. The first three states represent a triple dissociation of three domain-related streams: the dorsal stream of speech production, the ventral stream of semantic representation, and the central stream of phonology representation. The fourth state with weak connectivity strength serves as a baseline state, which may be related to non-verbal spontaneous thoughts (Kucyi, 2018; Kucyi, Esterman, Riley, & Valera, 2016; Kucyi, Hove, Esterman, Hutchison, & Valera, 2017). Such a framework endows the brain with a transient and dynamic synergy of functional-specific streams during language and speech processing (Hickok, 2022). The independence of State 1 endows the brain to flexibly and dynamically integrate it with State 2 or State 3. For example, for auditory sentence comprehension, which includes phoneme perception, phonological representation, word meaning retrieval, and meaning integration, hubs in states 1 and 3 will be recruited. For sentence repetition, which includes phoneme perception, phonological representation, and speech articulation, hubs in states 1 and 3 will be recruited. For sentence production, which includes word meaning retrieval, meaning integration, phonological representation, and speech articulation, hubs in states 1, 2, and 3 will be recruited (Binder, 2017).

The framework has also proven to have behavior and clinical relevance. In a large sample of 522 healthy subjects, we found that the four states significantly predicted individual linguistic performance (B. Yuan, Xie, Wang, et al., 2023). In 83 patients with left hemispheric gliomas involving language network, we found that the connectivity and topological properties of the four states were deficit-severity dependent and significantly predicated individual patients’ language scores (B. Yuan, Xie, Gong, et al., 2023). In this study, we found that the framework was susceptible to remote but acute lesions, showed severe network disruptions in the acute phase, and recovered in the chronic phase.

### The connectional diaschisis of basal ganglia and thalamus stroke to cortical language network dynamics

We found that the connectional diaschisis of cortical language network dynamics can be attributed to the cortico-subcortical dynamics. The four cortico-subcortical states we discovered were reliable in healthy subjects. The thalamus/basal ganglia appear to be always tightly connected, while the hippocampus and amygdala are always tightly connected, which forms two subnetworks of subcortical regions (Supplementary Figures 3 and 4). Favaretto et al. (2022) also identified similar connectivity patterns using a sliding window-based dynamic functional connectivity approach. The most probable reason for these tight connectivity patterns is their close anatomical proximity (Sepulcre et al., 2010) and functional communication (Shine et al., 2023). Additionally, in the study by Favaretto et al. (2022), they found no correlation between BG/thalamus and hippocampus/amygdala. However, in this study, we found that the two subnetworks exhibit positive connections at specific time points (State 2 in Supplementary Figures 3 and 4). In terms of dynamic interactions within cortical language network, these two subnetworks also differ. The connections between the thalamus/basal ganglia and the frontal language areas exhibit three modes: positive, negative, and weak connections, whereas connections between the STG and MTG are consistently antagonistic. Conversely, connections between the hippocampus/amygdala and the frontal language areas are consistently antagonistic, while they exhibit three modes of connection (positive, negative, and weak) with temporal lobe regions. Although the existing results cannot explain the functional relevance of the dynamic functional interactions between cortical and subcortical regions, the presence of such dynamic interactions is reasonable due to the close cortico-thalamo-cortical and cortico-striato-thalamic loops (Ranganath & Ritchey, 2012; Shine et al., 2023).

At a whole-brain level, the cortical language network we defined includes the classical language regions (IFG and STG), the auditory network, the temporal aspects of the default mode network, and the ventral aspects of the sensorimotor network (Supplementary Figures 1). Previous static functional connectivity (FC) studies consistently reveal functional connections between these networks and subcortical regions. For instance, Ji et al. (2019) identified close connections between the classical language network and specific subcortical areas (Supplementary Figures 1) and observed activation of these regions during language comprehension tasks, exhibiting a left lateralization pattern. Lu et al. (2021) indicated that the ventral aspects of the sensorimotor network are primarily associated with speech production. Neuroimaging studies have further shown co-activation of subcortical areas during various language and speech tasks (Turker et al., 2023). The default mode network is the most active network during rest, and subcortical regions closely connected to the cortical default network include the thalamus, hippocampus and caudate nucleus (Ji et al., 2019).

The cortico-subcortical dynamics in healthy subjects explain well why stroke in the thalamus/basal ganglia disrupts cortical language network dynamics. In the acute phase, the functional connectivity patterns of the first three states are severely disrupted, demonstrating the dual effects of hypoconnectivity and hyperconnectivity. The hubs in the first three states are dynamically correlated with the thalamus and basal ganglia. These observations suggested that acute injury to the thalamus/basal ganglia disrupts normal interactions with cortical language networks and mutual inhibition (Carson, 2020).

### The network normalization of cortical language network dynamics during language recovery

The restoration of functional connectivity and re-emergence of topological properties to sub-optimal patterns are crucial neural mechanisms for behavior and cognition recovery in lesioned brains, often referred to as the “network normalization” phenomenon (Otten et al., 2012; Ramsey et al., 2017; Siegel et al., 2018; Binke Yuan et al., 2022). However, previous studies on the“network normalization” phenomenon primarily relied on static functional connectivity analysis, overlooking the time-varying reconfiguration. Our findings suggest that language recovery is attributed to the normalization of cortical language network dynamics to a sub-optimal pattern, i.e., the cortical language dynamics conforms to a domain-segregation manner, but the topological properties cannot recover to optimal levels. It appears that sub-optimal cortical language dynamics are sufficient for language processing. In our recent study focused on patients with gliomas involving the left language areas, we found that the cortical language dynamics in patients without aphasia, attributed to neuroplasticity, were also sub-optimal (B. Yuan, Xie, Gong, et al., 2023).

There are limitations in this study. First, the subjects we selected have damage not only in the subcortical regions but also a considerable proportion of white matter damage (Figure 1). Undoubtedly, white matter disconnection is a critical cause (maybe the leading factor) of aberrant cortical language network dynamics (Griffis, Metcalf, Corbetta, & Shulman, 2019). However, it is difficult to distinguish the two potential factors in the current cases.

Second, while each subject underwent dense sampling (6–8 runs with at least 20 minutes of scanning), the sample size remained small. A larger sample size would be advantageous for lesion-deficits mapping analysis to depict the direct lesion effect to linguistic function. Also, a larger sample size endows with the opportunity to construct individual machine learning-based prediction modes (Siegel et al., 2016; B. Yuan, Xie, Gong, et al., 2023; B. Yuan et al., 2019), which would enhance the clinical relevance of these neuroimaging findings.

In conclusion, by adopting the dynamic meta-networking framework of language, we have illustrated the influence of acute lesion in basal ganglia and thalamus on cortical language network dynamics. Subcortical strokes impacted the domain-segregation cortical language network dynamics with dual effects. The restoration of cortical language network dynamics parallels the process of language recovery. These findings emphasize the critical role of cortico-subcortical interactions in language processing.

## Supporting information

SubcorticalStroke_dcc_supplementary_v2(1)

## Data Availability Statement

The stroke dataset has been publicly available at https://cnda.wustl.edu/data/projects/CCIR_00299. The DCC toolbox was available at t https://github.com/canlab/Lindquist_Dynamic_Correlation/tree/master/DCC_toolbox.

## References

Bell, P. T., & Shine, J. M. (2016). Subcortical contributions to large-scale network communication. Neurosci Biobehav Rev, 71, 313–322. doi:10.1016/j.neubiorev.2016.08.036

Binder, J. R. (2017). Current Controversies on Wernicke’s Area and its Role in Language. Curr Neurol Neurosci Rep, 17(8), 58. doi:10.1007/s11910-017-0764-8

Bonkhoff, A. K., Espinoza, F. A., Gazula, H., Vergara, V. M., Hensel, L., Michely, J., … Grefkes, C. (2020). Acute ischaemic stroke alters the brain’s preference for distinct dynamic connectivity states. Brain, 143(5), 1525–1540. doi:10.1093/brain/awaa101

Carson, R. G. (2020). Inter-hemispheric inhibition sculpts the output of neural circuits by co-opting the two cerebral hemispheres. J Physiol, 598(21), 4781–4802. doi:10.1113/JP279793

Choe, A. S., Nebel, M. B., Barber, A. D., Cohen, J. R., Xu, Y., Pekar, J. J., … Lindquist, M. A. (2017). Comparing test-retest reliability of dynamic functional connectivity methods. Neuroimage, 158, 155–175. doi:10.1016/j.neuroimage.2017.07.005

Copland, D. A., & Angwin, A. J. (2019). Subcortical contributions to language.

Copland, D. A., Brownsett, S., Iyer, K., & Angwin, A. J. (2021). Corticostriatal Regulation of Language Functions. Neuropsychology Review, 31(3), 472–494. doi:10.1007/s11065-021-09481-9

Corbetta, M., Ramsey, L., Callejas, A., Baldassarre, A., Hacker, C. D., Siegel, J. S., … Shulman, G. L. (2015). Common behavioral clusters and subcortical anatomy in stroke. Neuron, 85(5), 927–941. doi:10.1016/j.neuron.2015.02.027

Crosson, B. (2013). Thalamic mechanisms in language: a reconsideration based on recent findings and concepts. Brain Lang, 126(1), 73–88. doi:10.1016/j.bandl.2012.06.011

Damaraju, E., Allen, E. A., Belger, A., Ford, J. M., McEwen, S., Mathalon, D. H., … Calhoun, V. D. (2014). Dynamic functional connectivity analysis reveals transient states of dysconnectivity in schizophrenia. Neuroimage Clin, 5, 298–308. doi:10.1016/j.nicl.2014.07.003

Duffau, H., Moritz-Gasser, S., & Mandonnet, E. (2014). A re-examination of neural basis of language processing: proposal of a dynamic hodotopical model from data provided by brain stimulation mapping during picture naming. Brain Lang, 131, 1–10. doi:10.1016/j.bandl.2013.05.011

Fan, L., Li, H., Zhuo, J., Zhang, Y., Wang, J., Chen, L., … Jiang, T. (2016). The Human Brainnetome Atlas: A New Brain Atlas Based on Connectional Architecture. Cereb Cortex, 26(8), 3508–3526. doi:10.1093/cercor/bhw157

Favaretto, C., Allegra, M., Deco, G., Metcalf, N. V., Griffis, J. C., Shulman, G. L., … Corbetta, M. (2022). Subcortical-cortical dynamical states of the human brain and their breakdown in stroke. Nat Commun, 13(1), 5069. doi:10.1038/s41467-022-32304-1

Greene, D. J., Marek, S., Gordon, E. M., Siegel, J. S., Gratton, C., Laumann, T. O., … Dosenbach, N. U. F. (2020). Integrative and Network-Specific Connectivity of the Basal Ganglia and Thalamus Defined in Individuals. Neuron, 105(4), 742–758 e746. doi:10.1016/j.neuron.2019.11.012

Griffis, J. C., Metcalf, N. V., Corbetta, M., & Shulman, G. L. (2019). Structural Disconnections Explain Brain Network Dysfunction after Stroke. Cell Reports, 28(10), 2527–2540 e2529. doi:10.1016/j.celrep.2019.07.100

Herbet, G., & Duffau, H. (2020). Revisiting the Functional Anatomy of the Human Brain: Toward a Meta-Networking Theory of Cerebral Functions. Physiol Rev, 100(3), 1181–1228. doi:10.1152/physrev.00033.2019

Hickok, G. (2022). The dual stream model of speech and language processing. Handb Clin Neurol, 185, 57–69. doi:10.1016/B978-0-12-823384-9.00003-7

Hickok, G., & Poeppel, D. (2007). Opinion - The cortical organization of speech processing. Nature Reviews Neuroscience, 8(5), 393–402. doi:10.1038/nrn2113

Hodgson, V. J., Lambon Ralph, M. A., & Jackson, R. L. (2021). Multiple dimensions underlying the functional organization of the language network. Neuroimage, 241, 118444. doi:10.1016/j.neuroimage.2021.118444

Hwang, K., Shine, J. M., Bruss, J., Tranel, D., & Boes, A. (2021). Neuropsychological evidence of multi-domain network hubs in the human thalamus. Elife, 10. doi:10.7554/eLife.69480

Hwang, K., Shine, J. M., Cole, M. W., & Sorenson, E. (2022). Thalamocortical contributions to cognitive task activity. Elife, 11. doi:10.7554/eLife.81282

Ji, J. L., Spronk, M., Kulkarni, K., Repovs, G., Anticevic, A., & Cole, M. W. (2019). Mapping the human brain’s cortical-subcortical functional network organization. Neuroimage, 185, 35–57. doi:10.1016/j.neuroimage.2018.10.006

Kang, D. W., Chalela, J. A., Ezzeddine, M. A., & Warach, S. (2003). Association of ischemic lesion patterns on early diffusion-weighted imaging with TOAST stroke subtypes. Arch Neurol, 60(12), 1730–1734. doi:10.1001/archneur.60.12.1730

Kucyi, A. (2018). Just a thought: How mind-wandering is represented in dynamic brain connectivity. Neuroimage, 180(Pt B), 505–514. doi:10.1016/j.neuroimage.2017.07.001

Kucyi, A., Esterman, M., Riley, C. S., & Valera, E. M. (2016). Spontaneous default network activity reflects behavioral variability independent of mind-wandering. Proc Natl Acad Sci U S A, 113(48), 13899–13904. doi:10.1073/pnas.1611743113

Kucyi, A., Hove, M. J., Esterman, M., Hutchison, R. M., & Valera, E. M. (2017). Dynamic Brain Network Correlates of Spontaneous Fluctuations in Attention. Cerebral Cortex, 27(3), 1831–1840. doi:10.1093/cercor/bhw029

Lebo, M. J., & Box-Steffensmeier, J. M. (2008). Dynamic conditional correlations in political science. American Journal of Political Science, 52(3), 688–704.

Lindquist, M. A., Xu, Y., Nebel, M. B., & Caffo, B. S. (2014). Evaluating dynamic bivariate correlations in resting-state fMRI: a comparison study and a new approach. Neuroimage, 101, 531–546. doi:10.1016/j.neuroimage.2014.06.052

Lipkin, B., Tuckute, G., Affourtit, J., Small, H., Mineroff, Z., Kean, H., … Fedorenko, E. (2022). Probabilistic atlas for the language network based on precision fMRI data from >800 individuals. Sci Data, 9(1), 529. doi:10.1038/s41597-022-01645-3

Lu, J., Zhao, Z., Zhang, J., Wu, B., Zhu, Y., Chang, E. F., … Berger, M. S. (2021). Functional maps of direct electrical stimulation-induced speech arrest and anomia: a multicentre retrospective study. Brain. doi:10.1093/brain/awab125

Murphy, K., & Fox, M. D. (2017). Towards a consensus regarding global signal regression for resting state functional connectivity MRI. Neuroimage, 154, 169–173. doi:10.1016/j.neuroimage.2016.11.052

Nelson, B. G., Bassett, D. S., Camchong, J., Bullmore, E. T., & Lim, K. O. (2017). Comparison of large-scale human brain functional and anatomical networks in schizophrenia. Neuroimage Clin, 15, 439–448. doi:10.1016/j.nicl.2017.05.007

Otten, M. L., Mikell, C. B., Youngerman, B. E., Liston, C., Sisti, M. B., Bruce, J. N., … McKhann, G. M., 2nd. (2012). Motor deficits correlate with resting state motor network connectivity in patients with brain tumours. Brain, 135(Pt 4), 1017–1026. doi:10.1093/brain/aws041

Parsons, T. D., Rogers, S. A., Braaten, A. J., Woods, S. P., & Troster, A. I. (2006). Cognitive sequelae of subthalamic nucleus deep brain stimulation in Parkinson’s disease: a meta-analysis. Lancet Neurol, 5(7), 578–588. doi:10.1016/S1474-4422(06)70475-6

Radanovic, M., & Almeida, V. N. (2021). Subcortical aphasia. Current Neurology and Neuroscience Reports, 21, 1–15.

Radanovic, M., & Mansur, L. L. (2017). Aphasia in vascular lesions of the basal ganglia: A comprehensive review. Brain Lang, 173, 20–32. doi:10.1016/j.bandl.2017.05.003

Ramsey, L. E., Siegel, J. S., Baldassarre, A., Metcalf, N. V., Zinn, K., Shulman, G. L., & Corbetta, M. (2016). Normalization of network connectivity in hemispatial neglect recovery. Ann Neurol, 80(1), 127–141. doi:10.1002/ana.24690

Ramsey, L. E., Siegel, J. S., Lang, C. E., Strube, M., Shulman, G. L., & Corbetta, M. (2017). Behavioural clusters and predictors of performance during recovery from stroke. Nat Hum Behav, 1. doi:10.1038/s41562-016-0038

Ranganath, C., & Ritchey, M. (2012). Two cortical systems for memory-guided behaviour. Nat Rev Neurosci, 13(10), 713–726. doi:10.1038/nrn3338

Rangus, I., Fritsch, M., Endres, M., Udke, B., & Nolte, C. H. (2022). Frequency and phenotype of thalamic aphasia. Journal of Neurology, 269(1), 368–376. doi:10.1007/s00415-021-10640-4

Rubinov, M., & Sporns, O. (2010). Complex network measures of brain connectivity: uses and interpretations. Neuroimage, 52(3), 1059–1069. doi:10.1016/j.neuroimage.2009.10.003

Salvalaggio, A., De Filippo De Grazia, M., Zorzi, M., Thiebaut de Schotten, M., & Corbetta, M. (2020). Post-stroke deficit prediction from lesion and indirect structural and functional disconnection. Brain, 143(7), 2173–2188. doi:10.1093/brain/awaa156

Sepulcre, J., Liu, H., Talukdar, T., Martincorena, I., Yeo, B. T., & Buckner, R. L. (2010). The organization of local and distant functional connectivity in the human brain. PLoS Comput Biol, 6(6), e1000808. doi:10.1371/journal.pcbi.1000808

Shine, J. M., Lewis, L. D., Garrett, D. D., & Hwang, K. (2023). The impact of the human thalamus on brain-wide information processing. Nat Rev Neurosci, 24(7), 416–430. doi:10.1038/s41583-023-00701-0

Siegel, J. S., Ramsey, L. E., Snyder, A. Z., Metcalf, N. V., Chacko, R. V., Weinberger, K., … Corbetta, M. (2016). Disruptions of network connectivity predict impairment in multiple behavioral domains after stroke. Proc Natl Acad Sci U S A, 113(30), E4367–4376. doi:10.1073/pnas.1521083113

Siegel, J. S., Seitzman, B. A., Ramsey, L. E., Ortega, M., Gordon, E. M., Dosenbach, N. U. F., … Corbetta, M. (2018). Re-emergence of modular brain networks in stroke recovery. Cortex, 101, 44–59. doi:10.1016/j.cortex.2017.12.019

Silvestri, E., Moretto, M., Facchini, S., Castellaro, M., Anglani, M., Monai, E., … Corbetta, M. (2022). Widespread cortical functional disconnection in gliomas: an individual network mapping approach. Brain Commun, 4(2), fcac082. doi:10.1093/braincomms/fcac082

Thibault, S., Py, R., Gervasi, A. M., Salemme, R., Koun, E., Lovden, M., … Brozzoli, C. (2021). Tool use and language share syntactic processes and neural patterns in the basal ganglia. Science, 374(6569), eabe0874. doi:10.1126/science.abe0874

Thompson, W. H., Richter, C. G., Plaven-Sigray, P., & Fransson, P. (2018). Simulations to benchmark time-varying connectivity methods for fMRI. PLoS Comput Biol, 14(5), e1006196. doi:10.1371/journal.pcbi.1006196

Turker, S., Kuhnke, P., Eickhoff, S. B., Caspers, S., & Hartwigsen, G. (2023). Cortical, subcortical, and cerebellar contributions to language processing: A meta-analytic review of 403 neuroimaging experiments. Psychol Bull. doi:10.1037/bul0000403

Vigneau, M., Beaucousin, V., Herve, P. Y., Duffau, H., Crivello, F., Houde, O., … Tzourio-Mazoyer, N. (2006). Meta-analyzing left hemisphere language areas: phonology, semantics, and sentence processing. Neuroimage, 30(4), 1414–1432. doi:10.1016/j.neuroimage.2005.11.002

Vigneau, M., Beaucousin, V., Herve, P. Y., Jobard, G., Petit, L., Crivello, F., … Tzourio-Mazoyer, N. (2011). What is right-hemisphere contribution to phonological, lexico-semantic, and sentence processing? Insights from a meta-analysis. Neuroimage, 54(1), 577–593. doi:10.1016/j.neuroimage.2010.07.036

Wessels, T., Wessels, C., Ellsiepen, A., Reuter, I., Trittmacher, S., Stolz, E., & Jauss, M. (2006). Contribution of diffusion-weighted imaging in determination of stroke etiology. AJNR Am J Neuroradiol, 27(1), 35–39. Retrieved from https://www.ncbi.nlm.nih.gov/pubmed/16418352

Xie, H., Zheng, C. Y., Handwerker, D. A., Bandettini, P. A., Calhoun, V. D., Mitra, S., & Gonzalez-Castillo, J. (2019). Efficacy of different dynamic functional connectivity methods to capture cognitively relevant information. Neuroimage, 188, 502–514. doi:10.1016/j.neuroimage.2018.12.037

Xu, Y., Lin, Q., Han, Z., He, Y., & Bi, Y. (2016). Intrinsic functional network architecture of human semantic processing: Modules and hubs. Neuroimage, 132, 542–555. doi:10.1016/j.neuroimage.2016.03.004

Yuan, B., Fang, Y., Han, Z., Song, L., He, Y., & Bi, Y. (2017). Brain hubs in lesion models: Predicting functional network topology with lesion patterns in patients. Sci Rep, 7(1), 17908. doi:10.1038/s41598-017-17886-x

Yuan, B., Xie, H., Gong, F., Zhang, N., Xu, Y., Zhang, H., … Yan, J. (2023). Dynamic network reorganization underlying neuroplasticity: the deficits-severity-related language network dynamics in patients with left hemispheric gliomas involving language network. Cereb Cortex. doi:10.1093/cercor/bhad113

Yuan, B., Xie, H., Wang, Z., Xu, Y., Zhang, H., Liu, J., … Wu, J. (2023). The domain-separation language network dynamics in resting state support its flexible functional segregation and integration during language and speech processing. Neuroimage, 274, 120132. doi:10.1016/j.neuroimage.2023.120132

Yuan, B., Zhang, N., Gong, F., Wang, X., Yan, J., Lu, J., & Wu, J. (2022). Longitudinal assessment of network reorganizations and language recovery in postoperative patients with glioma. Brain Commun, 4(2), fcac046.

Yuan, B., Zhang, N., Yan, J., Cheng, J., Lu, J., & Wu, J. (2019). Resting-state functional connectivity predicts individual language impairment of patients with left hemispheric gliomas involving language network. Neuroimage Clin, 24, 102023. doi:10.1016/j.nicl.2019.102023

Zhao, Z., Liu, Y., Zhang, J., Lu, J., & Wu, J. (2021). Where is the speech production area? Evidence from direct cortical electrical stimulation mapping. Brain. doi:10.1093/brain/awab178

