## Supplementary material for "The connectional diaschisis and normalization of cortical language network dynamics after basal ganglia and thalamus stroke": SubcorticalStroke_dcc_supplementary_v2(1)

**Supplementary Table 1**

Supplementary Table 1 Anatomical regions，language-related behavioral domains, and paradigm classes of the language network. Coordinates are in the standard Montreal Neurologic Institute Space.

| ID | *x* | *y* | *z* | anatomy | Behavioral domains | Paradigm classes |
| --- | --- | --- | --- | --- | --- | --- |
| 1 | -4.6 | 15.7 | 53.2 | SFG | Execution. Speech, Phonology, Semantics, Speech | Word Generation (Covert and Overt) |
| 2 | 7.0 | 16.4 | 54.7 | SFG | Homolog of node 1 | - |
| 17 | -41.9 | 13.6 | 36.4 | MFG | Phonology, Semantics | Semantic. Monitor/Discrimination, Word Generation (Covert) |
| 18 | 42.2 | 11.9 | 38.4 | MFG | Homolog of node 17 | - |
| 29 | -45.6 | 13.0 | 23.4 | IFG | Phonology, Semantics, Speech, and Syntax | Phonological. Discrimination, Semantic. Monitor/Discrimination |
| 30 | 45.0 | 15.7 | 25.0 | IFG | Homolog of node 29 | - |
| 31 | -47.6 | 31.6 | 13.6 | IFG | Phonology, Semantics, Speech, and Syntax | Phonological. Discrimination, Semantic. Monitor/Discrimination, Word Generation (Covert and Overt) |
| 32 | 47.8 | 35.1 | 13.2 | IFG | Homolog of node 31 | - |
| 33 | -52.2 | 22.5 | 11.4 | IFG | Semantics, Speech, and Syntax | Reading (Covert), Semantic. Monitor/Discrimination, Word Generation (Covert and Overt) |
| 34 | 54.2 | 23.9 | 11.8 | IFG | Homolog of node 33 | - |
| 35 | -49.1 | 36.3 | -2.8 | IFG | Semantics, Speech, and Syntax | Semantic. Monitor/Discrimination, Word Generation (Covert) |
| 36 | 50.8 | 36.8 | -0.7 | IFG | Homolog of node 35 | - |
| 37 | -39.5 | 22.9 | 3.7 | IFG | Phonology, Semantics, Speech, and Syntax | Semantic. Monitor/Discrimination, Word Generation (Covert) |
| 38 | 42.1 | 22.0 | 3.2 | IFG | Homolog of node 37 | - |
| 39 | -51.3 | 13.2 | 6.0 | IFG | Phonology, Semantics, Speech | Music. Comprehension/Production, Recitation/Repetition. (Covert), Word Generation (Covert) |
| 40 | 53.6 | 14.3 | 11.8 | IFG | Homolog of node 39 | - |
| 53 | -49.5 | -7.1 | 38.8 | PrG | Execution. Speech | Reading (Overt), Recitation/Repetition. (Overt) |
| 54 | 54.9 | -2.0 | 33.3 | PrG | Execution. Speech | Reading (Overt), Recitation/Repetition. (Overt) |
| 63 | -49.1 | 4.7 | 30.5 | PrG | Orthography, Phonology, Semantics, Speech and Syntax | Phonological. Discrimination, Reading (Covert) |
| 64 | 51.1 | 7.2 | 30.9 | PrG | Homolog of node 63 | - |
| 71 | -53.7 | -32.1 | 12.4 | STG | Execution. Speech, Phonology, and Speech | Music. Comprehension/Production, Passive Listening, Phonological. Discrimination, Reading (Overt), Recitation/Repetition. (Covert and Overt) |
| 72 | 54.5 | -23.7 | 10.6 | STG | Execution. Speech, Phonology | Music. Comprehension/Production, Passive Listening, Phonological. Discrimination, Recitation/Repetition. (Overt) |
| 73 | -50.1 | -10.3 | 1.1 | STG | Execution. Speech, Phonology, Speech | Music. Comprehension/Production, Passive Listening, Phonological. Discrimination, Reading (Overt), Recitation/Repetition. (Overt) |
| 74 | 51.1 | -3.7 | -0.9 | STG | Execution. Speech, | Music. Comprehension/Production, Passive Listening, Recitation/Repetition. (Overt) |
| 75 | -62.8 | -32.8 | 7.4 | STG | Execution. Speech, Phonology, Semantics, Speech | Passive Listening, Phonological. Discrimination, Reading (Overt), Semantic. Monitor/Discrimination |
| 76 | 66.5 | -20.8 | 6.6 | STG | Execution. Speech, Phonology, Speech | Music. Comprehension/Production, Passive Listening, Phonological. Discrimination, Reading (Overt) |
| 77 | -45.1 | 10.5 | -19.4 | STG | Homolog of node 78 | - |
| 78 | 47.1 | 12.3 | -19.7 | STG | Speech | Film Viewing, Passive Listening |
| 79 | -55.1 | -3.2 | -10.1 | STG | Execution. Speech, Phonology, Semantics, Speech | Music. Comprehension/Production, Passive Listening, Phonological. Discrimination, Reading (Overt) |
| 80 | 55.8 | -12.5 | -5.2 | STG | Execution. Speech, Phonology, Semantics, Speech | Music. Comprehension/Production, Passive Listening, Phonological. Discrimination, Reading (Overt), Semantic. Monitor/Discrimination |
| 81 | -65.2 | -30.9 | -11.3 | MTG | Semantics | Semantic. Monitor/Discrimination |
| 82 | 64.5 | -29.2 | -13.2 | MTG | Homolog of node 81 | - |
| 83 | -53.2 | 2.2 | -29.6 | MTG | Language | Semantic. Monitor/Discrimination, Reading (Covert) |
| 84 | 51.1 | 5.7 | -31.8 | MTG | Language | Passive Listening |
| 85 | -58.9 | -57.6 | 4.3 | MTG | Semantics and Syntax | Semantic. Monitor/Discrimination, Word Generation (Overt) |
| 86 | 60.1 | -53.3 | 2.9 | MTG | Homologues of node 85 | Film Viewing |
| 87 | -58.5 | -19.8 | -9.4 | MTG | Phonology, Semantics, Speech, and Syntax | Passive Listening, Phonological. Discrimination, Reading (Covert), Semantic. Monitor/Discrimination |
| 88 | 58.3 | -15.4 | -10.1 | MTG | Semantics and Speech | Passive Listening, Phonological. Discrimination |
| 89 | -45.5 | -26.7 | -26.1 | ITG | Orthography and Semantics | Reading (Covert), Semantic. Monitor/Discrimination |
| 90 | 45.8 | -14.6 | -32.4 | ITG | Homolog of node 89 | - |
| 91 | -50.5 | -57.0 | -14.1 | ITG | Phonology and Semantics | Naming (Overt) |
| 92 | 53.5 | -52.4 | -18.5 | ITG | Phonology and Semantics | - |
| 93 | -43.7 | -2.9 | -41.4 | ITG | Semantics | Semantic. Monitor/Discrimination |
| 94 | 40.5 | -2.9 | -41.4 | ITG | Homolog of node 93 | - |
| 97 | -55.2 | -60.3 | -6.0 | ITG | Semantics and Speech | Film Viewing, Naming (Overt) |
| 98 | 54.2 | -56.9 | -8.6 | ITG | Homolog of node 97 | - |
| 99 | -58.8 | -42.1 | -16.0 | ITG | Orthography and Semantics | Naming (Overt), Reading (Covert), Semantic. Monitor/Discrimination, Word Generation (Overt) |
| 100 | 60.5 | -40.5 | -17.1 | ITG | Homolog of node 99 | - |
| 101 | -54.9 | -30.3 | -27.4 | ITG | Semantics | - |
| 102 | 53.7 | -30.3 | -26.3 | ITG | Homolog of node 101 | - |
| 103 | -32.4 | -16.6 | -32.3 | FuG | Semantics and Speech | Naming (Overt), Semantic. Monitor/Discrimination |
| 104 | 33.1 | -14.6 | -34.1 | FuG | Semantics | Naming (Overt), Semantic. Monitor/Discrimination |
| 105 | -30.6 | -64.4 | -14.1 | FuG | Orthography, Semantics, Speech | Naming (Covert and Overt) |
| 106 | 31.3 | -61.4 | -13.7 | FuG | Language | Naming (Covert and Overt) |
| 107 | -42.3 | -50.9 | -17.3 | FuG | Orthography, Phonology, Semantics, Speech | Naming (Covert and Overt), Phonological. Discrimination, Reading (Covert) and Semantic. Monitor/Discrimination |
| 108 | 42.7 | -49.1 | -18.6 | FuG | Orthography, Semantics | Naming (Covert) |
| 113 | -28.3 | -32.6 | -16.9 | PhG | Semantics | Naming (Overt), Semantic. Monitor/Discrimination |
| 114 | 28.8 | -30.7 | -17.5 | PhG | Homolog of node 113 | Passive Listening, Semantic. Monitor/Discrimination |
| 121 | -54.4 | -39.8 | 4.2 | pSTS | Phonology, Semantics, Speech, Syntax | Passive Listening, Phonological. Discrimination, Reading (Covert), Semantic. Monitor/Discrimination, Word Generation (Covert and Overt) |
| 122 | 52.9 | -36.8 | 3.1 | pSTS | Execution. Speech, Phonology, Semantics, and Speech | Passive Listening, Phonological. Discrimination, Reading (Overt) |
| 123 | -52.4 | -50.3 | 10.8 | pSTS | Orthography, Semantics, Speech, and Syntax | Reading (Covert), Semantic. Monitor/Discrimination |
| 124 | 56.5 | -40.1 | 12.5 | pSTS | Homologues of node 123 | Passive Listening |
| 143 | -46.8 | -64.7 | 25.8 | IPL | Language | Semantic. Monitor/Discrimination, |
| 144 | 53.0 | -54.1 | 24.4 | IPL | Homolog of node 143 | - |
| 155 | -50.4 | -15.8 | 42.1 | PoG | Execution. Speech | Recitation/Repetition. (Overt) |
| 156 | 50.3 | -14.2 | 43.7 | PoG | Execution. Speech | Reading (Overt), Recitation/Repetition. (Overt) |
| 157 | -55.8 | -14.0 | 16.2 | PoG | Execution. Speech | Recitation/Repetition. (Overt) |
| 158 | 55.2 | -10.2 | 15.0 | PoG | Execution. Speech | Recitation/Repetition. (Overt) |

*SFG*, superior frontal gurus; *MFG*, middle frontal gyrus; *IFG*, inferior frontal gyrus; *OrG*, orbital gyrus; *PrG*, precentral gyrus; *STG*, superior temporal gyrus; *MTG*, middle temporal gyrus; *ITG*, inferior temporal gyrus; *FuG*, fusiform gyrus; *PhG*, hippocampal gyrus; *pSTS*, posterior superior temporal sulcus; *IPL*, inferior parietal lobule. *PoG*, postcentral gyrus. The ID is the number of the parcel in the Brainnetome atlas (Fan et al., 2016). Here we only summarized the language-related behavioral domains and paradigm classes; the full behavioral domains and paradigm classes for each node are available at <http://atlas.brainnetome.org/bnatlas.php>.

**Supplementary Figures**

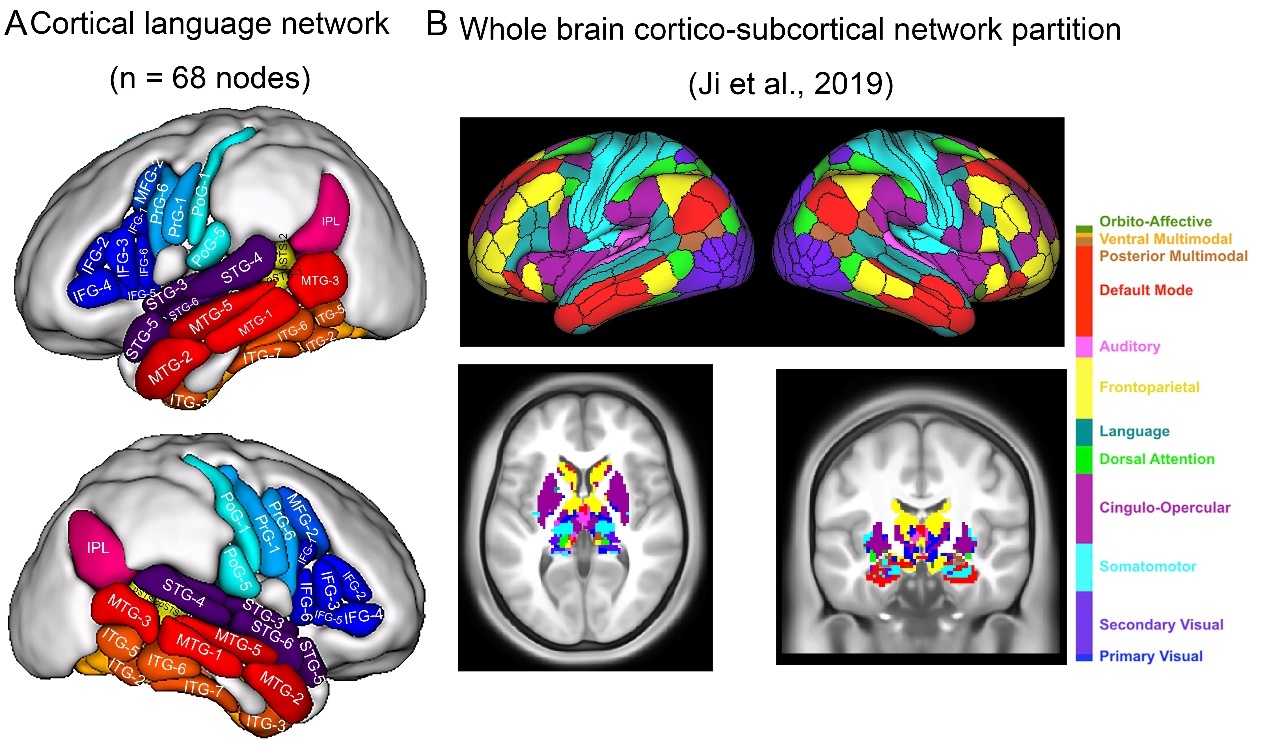

**Supplementary Figure 1.** The cortical language network and the 11 large-scale brain networks. A: 68 cortical regions involved in language processing were selected from the Brainnetome atlas (Fan et al., 2016). B: To delineate the resting-state brain networks encompassing the cortical language network we defined, we have redrawn the 12 whole-brain resting-state networks based on functional connectivity as defined by Ji et al. (2019). It can be observed that the language network we defined includes the typical language network (IFG and STG), auditory network, the default mode network, and the ventral part of the somatomotor network. These networks each have corresponding regions in the subcortical areas.

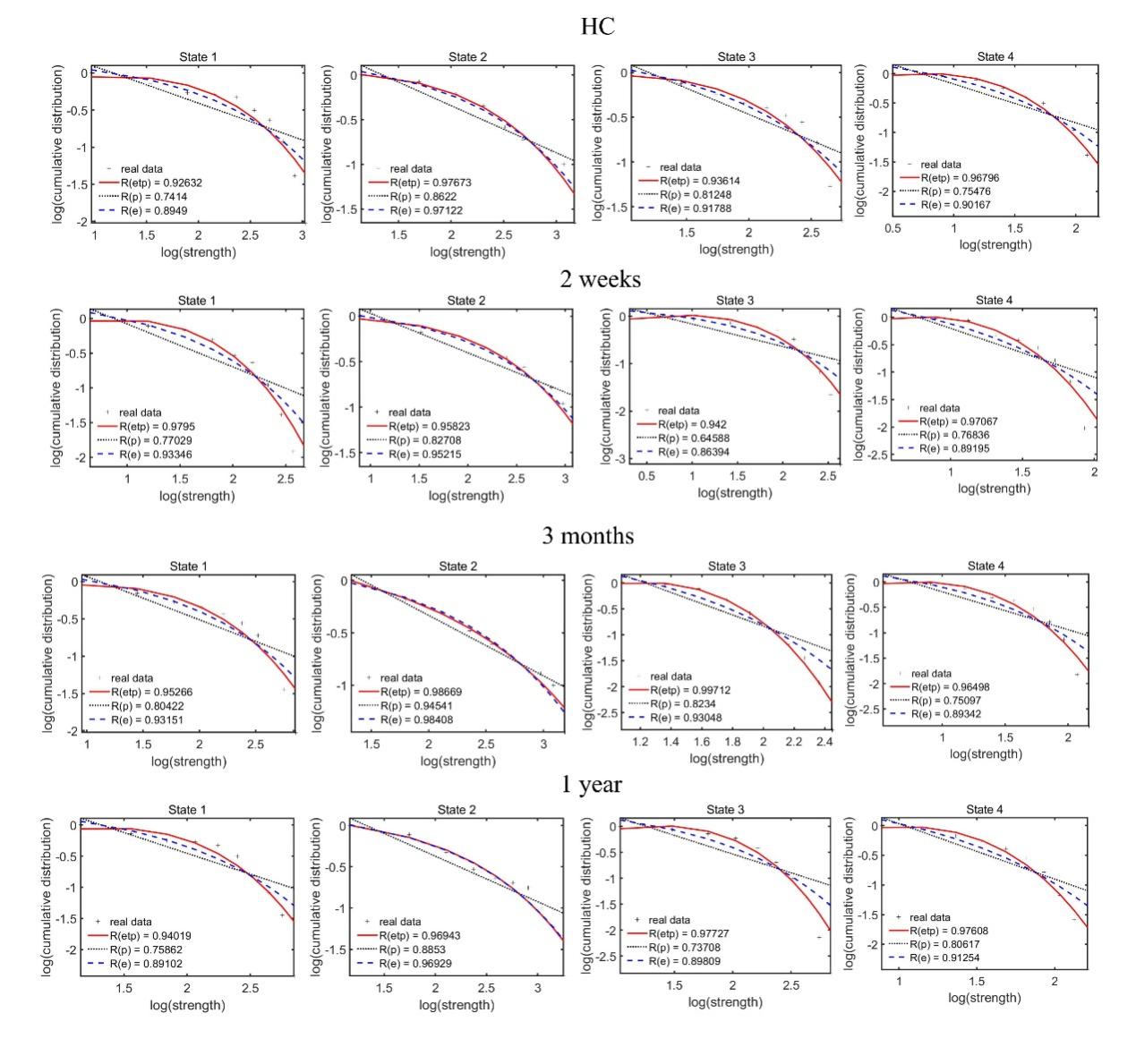

**Supplementary Figure 2.** The log-log plots of the cumulative nodal strength distributions. The plus sign (black) represents observed data, the solid line (red) is the fit of the exponentially truncated power-law, $P \left( x \right) \sim x^{\alpha-1}exp(\frac{x}{x_{c}})$, the dashed line (blue) is an exponential, $P \left( x \right) \sim\exp\left( \frac{x}{x_{c}} \right)$, and the dotted line (black) is a power-law, $P \left( x \right) \sim x^{\alpha-1}.$*R*^2^ was calculated to assess the goodness-of-fit. A larger value indicates a better fitting: *R_etp_*, *R^2^* for the exponentially truncated power-law; *R_e_*, *R^2^* for the exponential; *R_p_*, *R^2^* for the power-law fit. The exponentially truncated power-law is the best fitting for all 4 states, which suggest a long-tailed broad-scale topologies and a large proportion of network connectivity will be concentrated on a subset node (i.e., hubs).

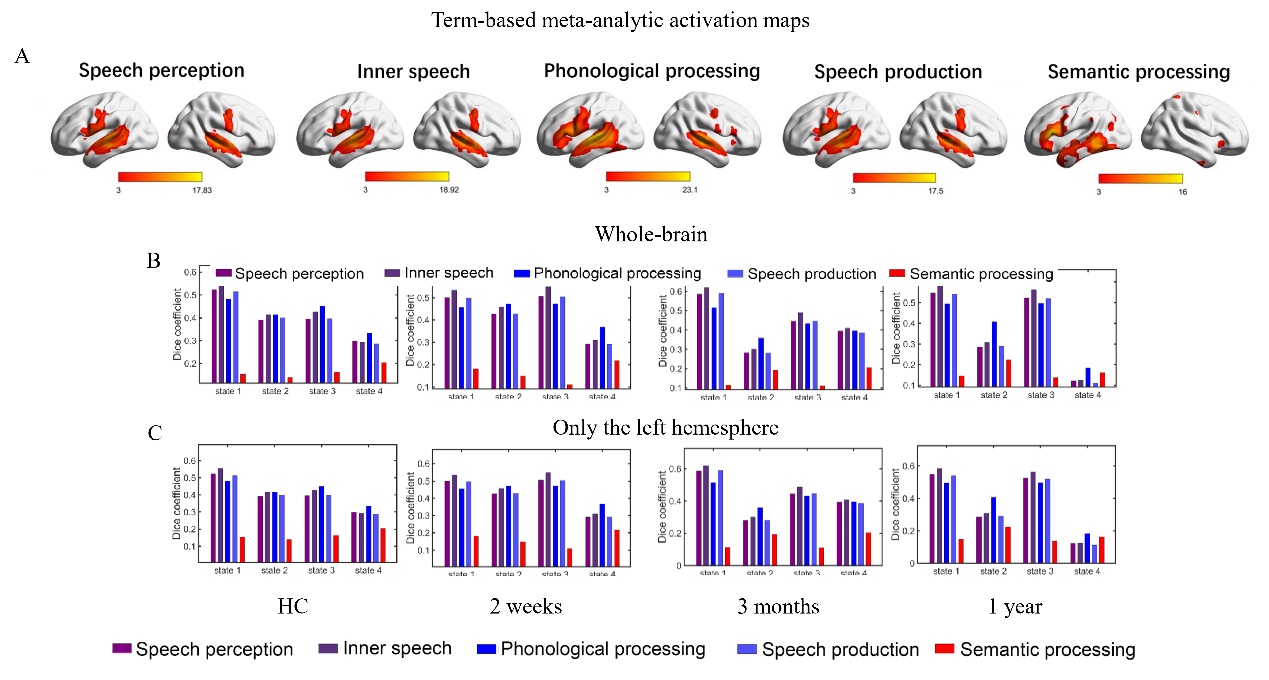

**Supplementary Figure 3.** Functional relevance of hub distributions. A. The meta results of speech perception, inner speech, phonological processing, speech production, and semantic processing were from 'NeuroQuery’ (https://neuroquery.org/) (Dockes et al., 2020). Each map was thresholded at Z = 3 (a typical value used by NeuroQuery) for illustrative purposes and only positive results were shown. B&C: the dice coefficients between binary images of hub nodes and meta results. Considering the left-lateralized activations of meta results, the dice coefficients were calculated at the whole-brain level and in the left hemisphere.

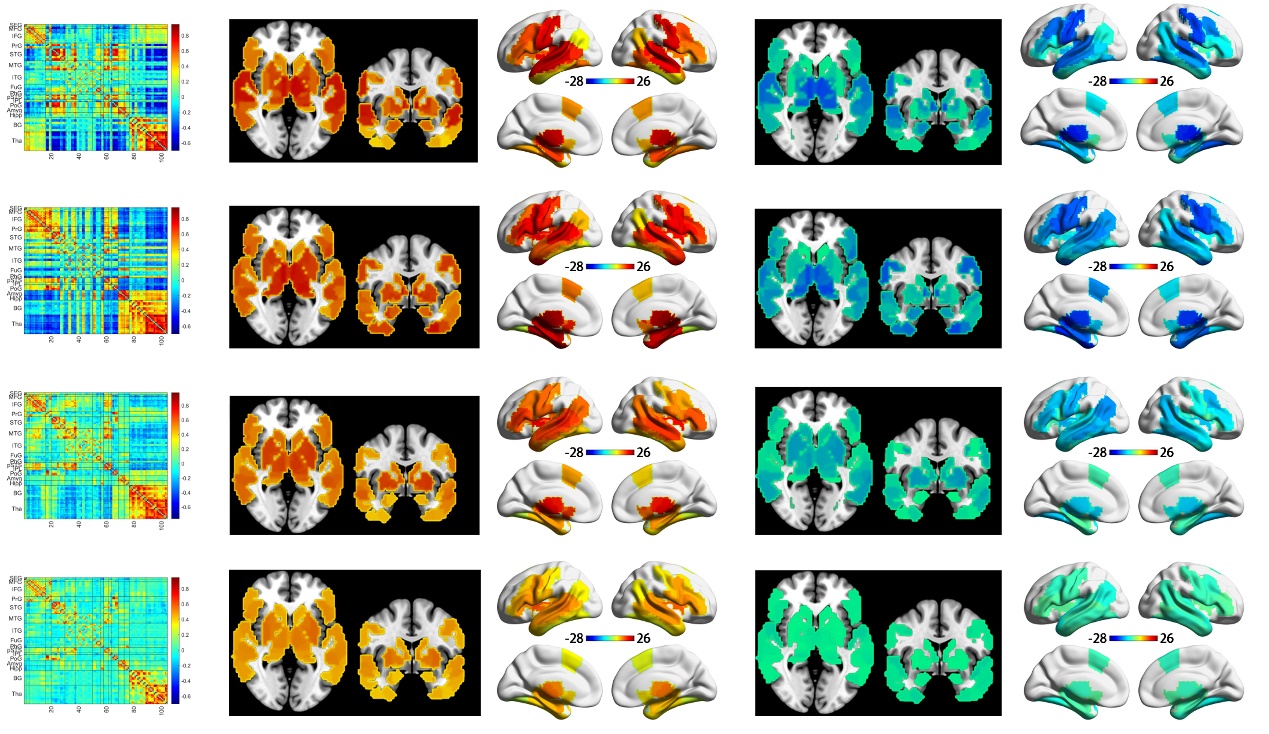

**Supplementary Figure 4.** The dynamic functional connectivity between cortical language areas and subcortical areas of HCs (n = 25). Figures in the left column show the four temporal-reoccurring DFC states. Figures in the right four columns show the nodal strengths of positive and negative connections in volumetric and surface space.

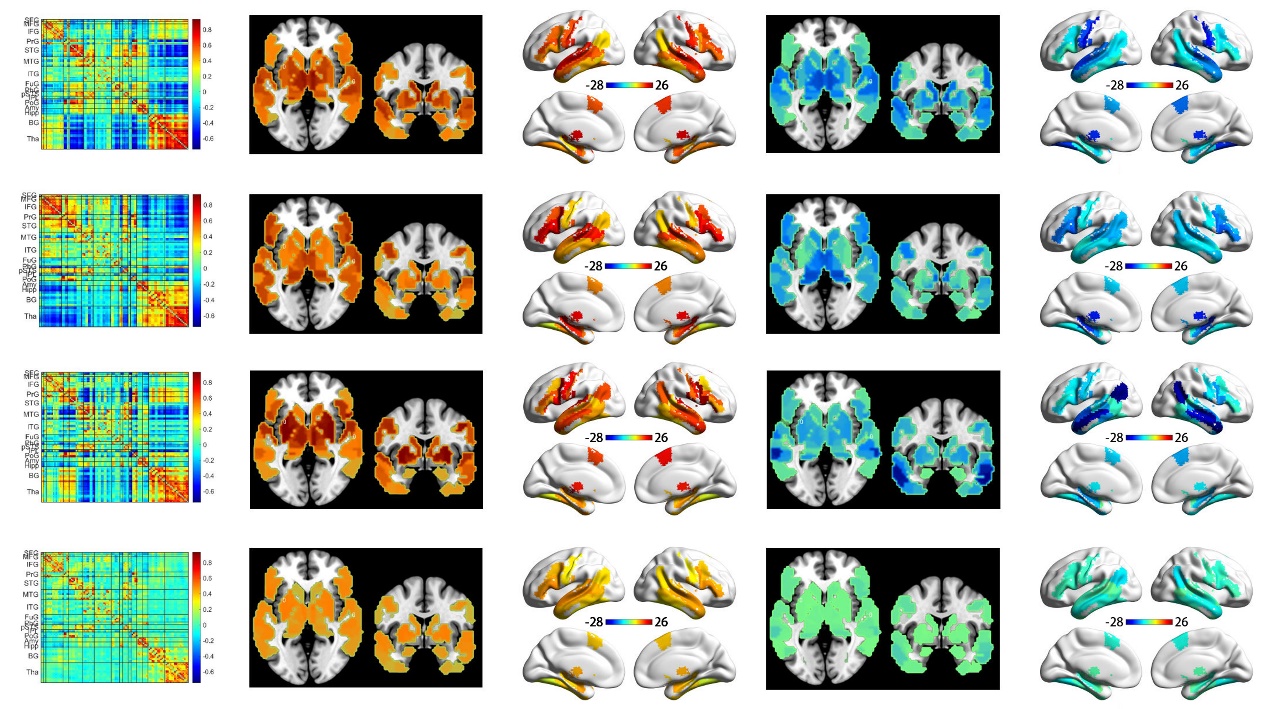

**Supplementary Figure 5.** The cortico-subcortical dynamics in young healthy subjects (n = 192).

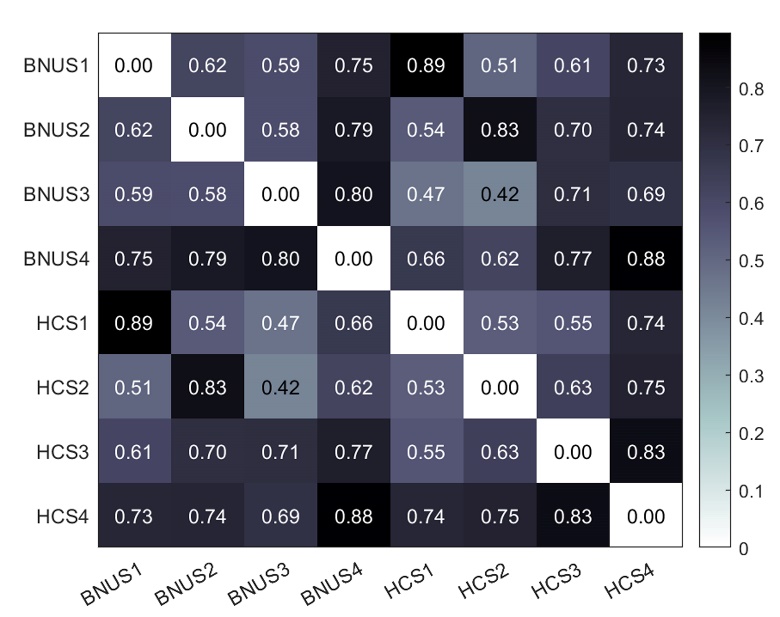

**Supplementary Figure 6.** The spatial correlation of the four states of cortico-subcortical dynamics in HCs (n = 25) and young healthy subjects (n = 192).

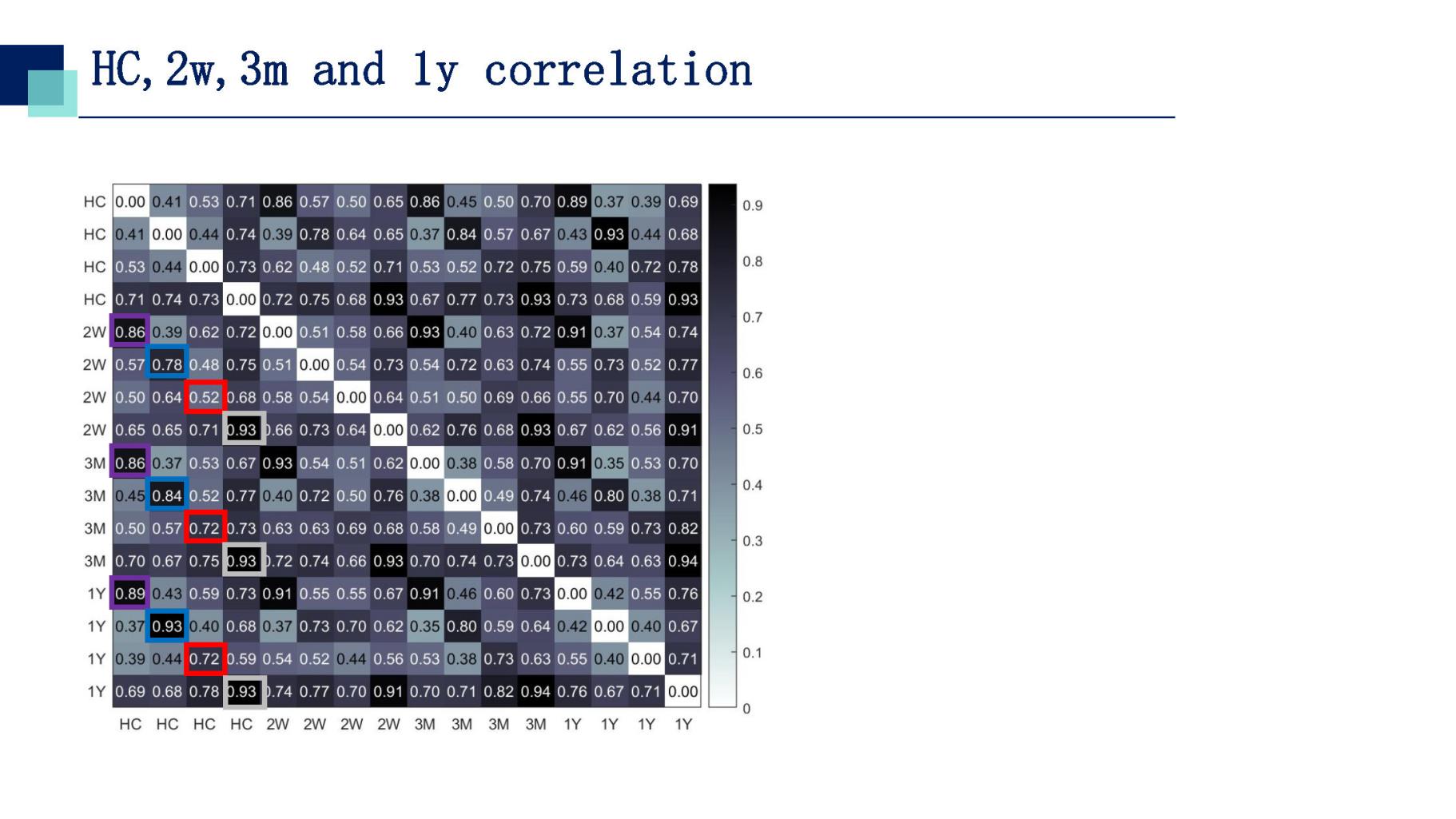

**Supplementary Figure 7.** The spatial correlation coefficients among the four states of HCs and patients. During language recovery, an increase in spatial similarity between patients and HCs was observed, especially for states 2 and 3.

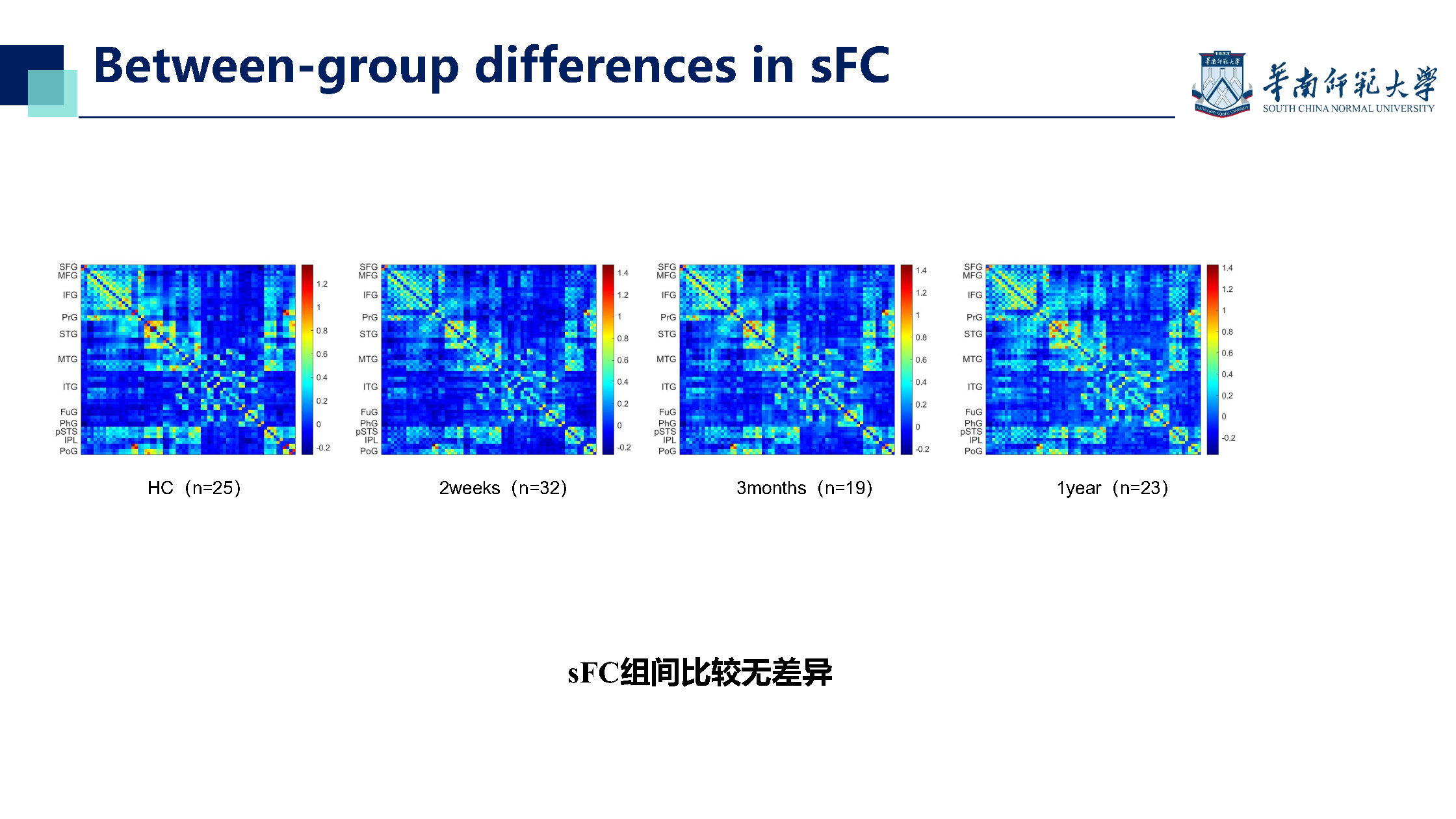

**Supplementary Figure 8.** The mean static functional connectivity of HCs and patients after stroke.
